# Reshaping the *Hexagone*: the genetic landscape of modern France

**DOI:** 10.1101/718098

**Authors:** Simone Andrea Biagini, Eva Ramos-Luis, David Comas, Francesc Calafell

**Author notes:** Address correspondence to Francesc Calafell, Departament de Ciències Experimentals i de la Salut, Institute of Evolutionary Biology (CSIC-UPF), Universitat Pompeu Fabra, Carrer Doctor Aiguader 88, 08005 Barcelona, Catalonia, Spain.

## Abstract

Unlike other European countries, the human population genetics and demographic history of Metropolitan France is surprisingly understudied. In this work, we combined newly genotyped samples from various zones in France with publicly available data and applied both allele frequency and haplotype-based methods in order to describe the internal structure of this country, by using genome-wide single nucleotide polymorphism (SNP) array genotypes. We found out that French Basques are genetically distinct from all other populations in the *Hexagone* and that the populations from southwest France (namely the Gascony region) share a large proportion of their ancestry with Basques. Otherwise, the genetic makeup of the French population is relatively homogeneous and mostly related to Southern and Central European groups. However, a fine-grained, haplotype-based analysis revealed that Bretons slightly separated from the rest of the groups, due mostly to gene flow from the British Isles in a time frame that coincides both historically attested Celtic population movements to this area between the 3th and the 9th centuries CE, but also with a more ancient genetic continuity between Brittany and the British Isles related to the shared drift with hunter-gatherer populations. Haplotype-based methods also unveiled subtle internal structures and connections with the surrounding modern populations, particularly in the periphery of the *Hexagone*.

## Introduction

Located in the center of Western Europe, Metropolitan France has historically acted as a bridge connecting Northern Europe to the Mediterranean and the Iberian spaces. The geographical position of France strongly affected the history of the settlement of the different parts of the territory, whose continuous fragmentation through time is attested by the large number of populations and cultures that settled this area. Greeks, Romans and Celtic tribes from central Europe shaped a first internal structure between the 6th and the 1st centuries BCE, while waves of barbarian invasions (Alamanni, Burgundians, Visigoths, Franks, and Celts) strongly impacted the population landscape of France during the 5th century CE^1^. During the 9th and 10th centuries CE, foreign invasions from all sides also influenced the territory: Muslims and Saracens from North Africa coming through Iberia, Hungarian Magyar from the east, and Vikings (*Northmen*) from the north^1^. Nowadays, France is a cosmopolitan country whose society is shaped by a plurality of lifestyles and truly different ethno-cultural diversity. Without any doubt, the impact of political refugees throughout the 20th century, or of the immigration from colonized countries to mainland France, such as the migration of Arabs and Berbers from Algeria which was the most extensive of all colonial migrations to Western Europe before the 1960s ^2^, enriched the modern genetic landscape of the French territory. However, it is beyond our intention to explore this plethora of recent genetic contributions here, which can be quantified much more precisely with demographic analyses. Instead, we can apply genomic tools to excavate a deeper and ancient genetic background.

At the light of this complex past, the genetic landscape of France has been poorly analyzed, especially in recent times. The first studies with classical markers defined a general heterogeneous pattern considering different geographical arrangements such as military districts, historical provinces, and regions^3,4^. With his synthetic maps, Cavalli-Sforza proposed that this heterogeneity was a consequence of differential Neolithic influences between northern and southern France, and also pointed out a differentiation for Brittany and Gascony ^5^. More recently, studies on mitochondrial DNA highlighted a general homogeneity when the samples were distributed among the 22 regions established in 1982 and historic provinces ^6,7^. Generally, the mtDNA haplogroup composition of French people did not differentiate neither internally, nor from the surrounding European genetic landscape ^6,7^. On a microgeographical scale, Brittany showed affinity with Scandinavia and Britain, while French Basques stood out for a high frequency of haplogroup H, suggesting a link with the Neolithic diffusion in Europe ^6,7^. In agreement with the homogeneity described by mtDNA studies, the Y-chromosome diversity strongly pointed out a lack of differentiation between the distinct groups when samples were organized on a regional scale. Even in this case, Brittany represented an exception, showing a lower Y-chromosome diversity that was interpreted as consequence of a possible founder effect, plus an isolation process ^8^. Based on autosomal variants, a genome-wide study on Western France did not find any differentiation among the distinct groups organized on a regional geographical distribution ^9^. Even in this case, the only outlier was Brittany, whose higher linkage disequilibrium suggested a lower effective population size, thus supporting the hypothesis of isolation inferred by the outcomes of the Y-chromosome analyses. Furthermore, in agreement with mitochondrial studies, Bretons were found to be admixed with individuals from the British Isles ^9^. In this work, we present a comprehensive genome-wide study on France, using both allele frequency and haplotype-based methods, to determine the minimal meaningful geographic unit of genetic differentiation within France, describe the geogenetical landscape patterns within France, and trace the historic and ancient sources of gene flow into the *Hexagone*.

## Material and Methods

### Dataset arrangement and genotypes

In this study, informed consent was obtained from 331 individuals from different French departments. Internal Review Board approval for this work was granted by CEIC-PSMAR ref. 2016/6723/I. These samples were compiled by the Institute of Forensic Sciences, University of Santiago de Compostela, and most of them were first reported in an analysis of Y-chromosome markers in ref. ^8^. As specified in the latter work, all the subjects and their parents were born in mainland France and bore a French surname. DNA was extracted from blood samples as described in Ramos-Luis *et al.* ^8^. A total of four Axiom ® Genome-Wide Human Origins Arrays (∼629 K SNPs) ^10^ were genotyped at the Centro Nacional de Genotipado - Universidade de Santiago de Compostela facility. Genotype calling was performed running four different batches according to the Affymetrix Best Practices Workflow implemented in the software Axiom™ Analysis Suite 2.0. Out of 331 samples, 52 failed the genotyping process and a total of 279 samples were retained. Three additional samples were removed following an Identity-by-descent analysis (IBD) since they displayed a Proportion IBD value ≥ 0.125 (minimum threshold for removing relatedness equal or higher than a third degree). Eventually, 276 samples were retained. To complete the French dataset, 79 additional samples from a public source ^11^ and 60 from unpublished data (from an ongoing study on the Basque Country and the Franco-Cantabrian region; samples are subset from those in ref. ^12^) were added to the original 276, leading to a total of 415 samples. In a preliminary part of this work, 20 out of the 276 samples were identified as outliers and removed from the study (see Supplementary Figure 1 and caption). Thus, the complete dataset included 256 newly genotyped samples, plus 139 additional ones, for a final group of 395 samples (Dataset A) distributed among 20 different French departments (see Supplementary Figure 2 for the geographical distribution). For the allele frequency analyses, as comparison with external populations, a total of 333 samples were added to Dataset A, forming Dataset B. This external group included 218 samples among Germany, Norway, Spain, Italy, England, Ireland, and Scotland ^11^, together with 107 samples from the Spanish autonomous communities of Catalonia, Valencian Community, and Balearic islands ^13^, and 8 additional samples from South Italy (Naples) newly genotyped with Axiom ® Genome-Wide Human Origins Arrays (∼629 K SNPs). Further 799 samples from external populations ^11^ were added to the previous ones when applying haplotype-based methods (Dataset C). Lastly, in the analysis with ancient data, 282 ancient samples ^11^ were added to the previous dataset, with the only exclusion of the 122 sub-Saharan African samples (Dataset D) since their presence would have reduced the resolution for the distribution of the rest of the samples in the PCA, masking signals of admixture in the dedicated analyses (see Supplementary Figure 3 for the geographical distribution of the modern samples from Datasets B and C, and Supplementary table 1 for a summary of the different dataset composition).

### Data Quality Control

Data were prepared using PLINK1.9 ^14^. Uniparental markers and X-chromosome variants were excluded. For the French dataset, a preliminary set of filters were applied to each group separately before the merging process. We filtered out all variants with missing call rates greater than 5%, those that failed Hardy-Weinberg test at p < 10 ^−5^, and samples with more than 10% missing genotype data. After merging, only variants common to the three datasets were retained and SNPs with a minor allele frequency (MAF) below 5% were excluded, resulting in a final 343,884 variants used for haplotype-based methods (Dataset A). For the analyses that needed a set of independent markers, SNPs were pruned setting a pairwise linkage disequilibrium maximum threshold of 0.5, a window of size 200 and a shift step of 25. Eventually, the pruned data retained 142,803 variants (Dataset A). In the analyses that included the external populations, only the pruned dataset, consisting in 154,889 SNPs, was used for the allele frequency analyses (Dataset B), while a set of 380,697 variants was retained in the haplotype-based methods (Dataset C). Regarding Dataset D, a set of 163,631 SNPs was retrieved after pruning (See Supplementary table 1 for a summary).

### Statistical analyses

Eigenvectors were computed using the SmartPCA program in Eigenstrat software package (v. 13050) ^15^. For Dataset D, we used the option lsqproject:YES when projecting ancient on top of the modern samples. Results were plotted in R (v 3.0.1).

The *F_ST_* fixation index was computed using the SmartPCA tool (v. 13050) from the Eigenstrat software package. Results were produced in Rstudio ^16^ using R version 3.4.4 ^17^. The *F_ST_* matrix was used together with a geographic distance matrix produced with The Geographic Distance Matrix Generator (v. 1.2.3, available from http://biodiversityinformatics.amnh.org/open_source/gdmg) in order to perform a Mantel test correlation using the ade4 ^18^ library in R. Results were displayed using ggplot2 ^19^ and reshape ^20^ libraries.

Based on different hierarchical levels (within Departments, Between Departments within Areas/Regions, Between Areas/Regions; see Supplementary Figure 4 for a visual representation of the used Areas and Regions), AMOVA was performed using the *poppr.amova* function in R package poppr (v. 2.8.1) ^21,22^ and significance was tested with the *randtest* function implemented in R package ade4. For every percentage of variance, a p-value was calculated based on 1000 permutations.

Patterns of population structure were explored, in both Dataset B and D, using ADMIXTURE ^23^ testing from K=2 to K=10 ancestral clusters and using 10 independent random seeds. Results were represented using the software pong ^24^. For Dataset B, admixture was formally tested with f3 statistics computed using the *qp3Pop* function implemented in Admixtools ^10^, while outgroup-f3 statistics were tested for Dataset D in the form of f3(Ancient, X; Mbuti), where 3 Mbuti samples from ref. ^11^ were added to Dataset D (1690 total samples, same variants as in Dataset D).

### EEMS (Estimated Effective Migration Surface)

EEMS ^25^ analysis was run using Dataset A (142,803 variants from the pruned file). With a matrix of average pairwise genetic dissimilarities calculated using the internal program bed2diffs, a sample coordinates file, and a habitat coordinates file generated using Google Earth Pro (v. 7.3.2.5495), we performed 10 pilot runs of 6 million MCMC iterations each, with 3 million burn-in, and a thinning interval of 30,000. A second set of 5 runs was then performed restarting the chain with the highest likelihood with 4 million MCMC iterations, 1 million burn-in, and thinning interval of 10,000. The density of the population grid was set to 300 demes, and random seeds were used for each one of the runs. We used the default hyperparameter values but tuned some of the proposed variances to improve convergence in the second set of runs. Results for the chain with the highest likelihood were displayed using eems.plots function in the R package rEEMSplots.

### Haplotype-based analysis

Two different analyses were performed: one on the internal French population only (Dataset A), and one also including external populations (Dataset C). In both cases, phasing was performed using the software Shapeit (v. v2.r837) ^26,27^. When running ChromoPainter ^28^, all samples were used as both recipients and donors, ^28^without any population specification (-a option) and not allowing self-copying. First, the parameters for the switch rate and global mutation probability were estimated with the EM algorithm implemented in ChromoPainter using the parameters -i 15 -in -iM for chromosomes 1, 7, 14, and 20 for all the samples. This step allows to estimate the two parameters that will be then averaged for all chromosomes. The outcome for the average weighted values for the global mutation probability and the switch rate parameters were respectively 0.000745 and 266.67196 for Dataset A, and 0.000586 and 237.50784 for Dataset C. In a second step, ChromoPainter was run for all chromosomes using the two fixed parameters. Later, the final coancestry matrices for each chromosome were combined using the tool Chromocombine. The latter also estimates the C parameter which is needed for the normalization of the coancestry matrix data when we run fineSTRUCTURE in order to identify the population structure. The MCMC of fineSTRUCTURE was run using 1000000 burn-in iterations (flag -x), 2000000 iterations sampled (flag -y), and thinning interval of 10000 (flag -z). Eventually, the fineSTRUCTURE tree was estimated running three different seeds and using the flags - X -Y -m T that allow to build the sample relationship tree. In the analysis on Dataset C, the work was then divided in two phases. In the first one, ChromoPainter and fineSTRUCTURE were rerun, this time silencing France in order to define the external groups only. In the second phase, fineSTRUCTURE was rerun using the “force file” option (-F), using “continents” as donor groups (represented by the external groups defined in the first phase); -F is a function that allows to exclude the donor representation in the building tree phase and focus on the distribution of the recipient groups, represented by the French samples only. We then applied the non-negative-least-squares (nnls) function from GLOBETROTTER ^29^ in order to describe the ancestry profiles for the French groups we detected with the “force file” option. We then used GLOBETROTTER in order to describe admixture events, sources and dates. More details about the usage of GLOBETROTTER are reported in Supplementary note 1.

## Results

### Internal genetic structure in France

In order to define the best geographical partitioning of genetic differentiation, a hierarchical analysis of molecular variance (AMOVA) was performed with areas or regions as major grouping factors. We determined first the proportion of genetic variation partitioned among geographic areas, among departments within geographic areas, and within departments. We next tested the proportion of genetic variation partitioned among regions (considering the 13 regions established in 2016), among departments within regions, and within departments. A further AMOVA was performed only testing the proportion of genetic variation partitioned among and within departments. As shown in Table 1, in all cases the main contribution to the genetic variance was found at the lowest hierarchical level (variation within departments), while differences among regions resulted in a negative value that could be interpreted as zero, meaning absence of any structure at this level. Conversely, differences among areas displayed positive values, supporting the role of areas as more reliable grouping factors of genetic variations when considering wider sample distributions. Finally, the results for the variation between departments, also supported by significant p-values in all the AMOVA analyses, pointed to the fact that this level of stratification might be a better representation for the minimal unit of genetic differentiation. Based on these results, samples were distributed on the map according to the departmental locations (Supplementary Figure 2) and all the subsequent analyses considered this grouping factor, although, given their known cultural and genetic identity, we retained Basque-speakers as a separate group in the Pyrénéés-Atlantiques department. A first Principal Component Analysis (PCA) showed two distinct groups separated along the first PC (Figure 1A): the Basque samples on the right part of the plot, against most of the rest of the samples on the left one, within which a structure cannot be defined. These two major groups are connected by a “bridge” of samples represented by non-Basque*-* speaking individuals from the Gascony region in the southwestern corner of France. When we averaged the eigenvalues for the first two PCs and represented the same PCA, together with standard deviation (SD) values for each group, no evident pattern could still be discerned beyond the separation of Basques and Gascons (Supplementary Figure 5A). When we removed both Basque and Gascon samples from the analysis (Figure 1B), the resulting PCA showed some internal pattern of differentiation, more clearly defined by the average PCA (Supplementary Figure 5B), in which samples from the departments belonging to the northwestern region of Brittany seem to form a cluster on the left part of the plot.

**Figure 1.**
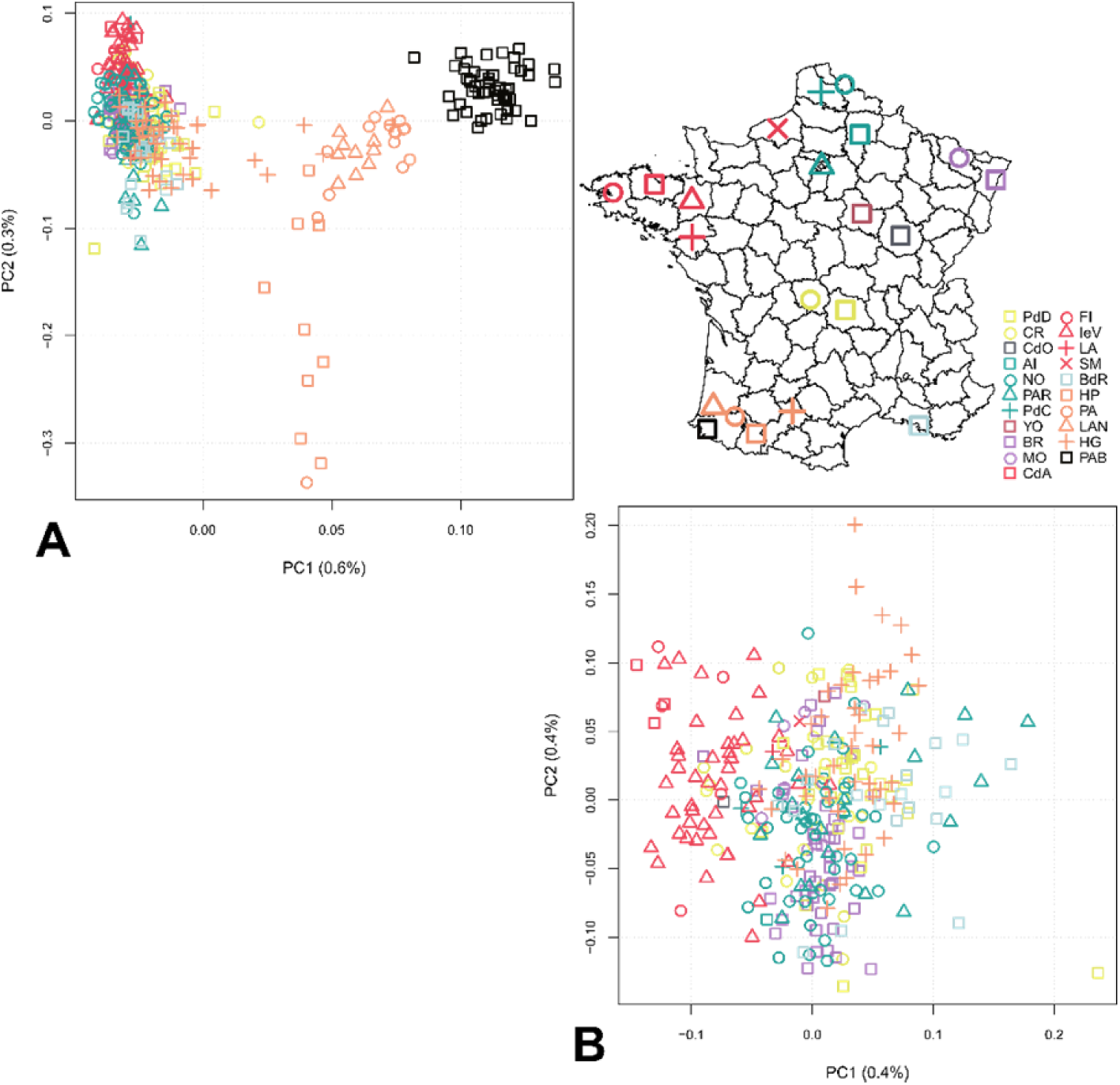
Principal Component Analysis of French samples (dataset A) with **A)** Basque and Gascon samples, and **B)** without them. Colors correspond to distinct geographic areas, while different symbols with the same color represent distinct departments in each area (See map distribution). However, Basques are colored differently than the non-Basque-speaking samples from that same area, but symbols recall the departments they share with the non-Basque-speaking groups.

**Table 1.** Hierarchical analysis of molecular variance (AMOVA). Results for percentage of total variance, Ф-statistics, and p-values are reported for the three distinct analyses. A) proportion of genetic variation partitioned among geographic areas, among departments within geographic areas, and within departments; B) proportion of genetic variation partitioned among regions, among departments within regions, and within departments; C) proportion of genetic variation partitioned among departments and within departments

| | Groupings | % Total variance | Φ-statistics | p |
| --- | --- | --- | --- | --- |
| **A** | Variations Between Areas | 0.02 | $\Phi_{ST} = 0.0026$ | 0.4605 |
| | Variations Between Departments Within Areas | 0.23 | $\Phi_{ST} = 0.0023$ | 0.0009 |
| | Variations Within Departments | 99.73 | $\Phi_{ST} = 0.00027$ | 0.0009 |
| **B** | Variations Between Regions | -0.054 | $\Phi_{ST} = -0.0005$ | 0.7362 |
| | Variations Between Departments Within Regions | 0.3 | $\Phi_{ST} = 0.003$ | 0.0009 |
| | Variations Within Departments | 99.74 | $\Phi_{ST} = 0.0025$ | 0.0009 |
| **C** | Variations Between Departments | 0.26 | $\Phi_{ST} = 0.0026$ | 0.0009 |
|  | Variations Within Departments | 99.73 |  |  |

### Patterns of gene flow within France

In the genetic variation computed with the *F_ST_* analysis, a general homogeneous pattern was found, with fine scale values of differentiation between some departments. The southwestern samples (Basques and Gascons) showed the highest values of differentiation with the northwestern departments reaching scores between 0.008 and 0.009 for the Basque-speaking samples, and between 0.004 and 0.006 for the non-Basque-speaking ones (Supplementary Figure 6A, left), followed by lower values of differentiation with the northern and northeastern departments. Without the southwestern samples, the main differentiation was recorded between the northwestern departments and the southeastern corner of the country, with a highest value of differentiation around 0.002 between the southeastern department of Bouches-du-Rhône (BdR) and the northwestern Breton department of Côtes-d’Armor (CdA) (Supplementary Figure 6B, left). Lower levels of differentiation were locally found among the departments in the northwest, and among those in the north together with the northeastern ones. A Multidimensional Scaling analysis (MDS) based on the *F_ST_* matrices clearly showed how the southwestern samples separate from the rest of the groups (Supplementary Figure 6A, right), and how the Breton departments do the same once the Gascon and Basque samples are removed (Supplementary Figure 6B, right). A Mantel test of isolation by distance (IBD) between the *F_ST_* values and the geographical distances showed a positive and statistically supported correlation (R^2^=0.332, P=0.001) (Supplementary Figure 7A), moving to even more positive values when the southwestern samples were removed (R^2^=0.432, P=0.001) (Supplementary Figure 7B). Next, we used the EEMS analysis, a method for visualizing genetic diversity patterns, and found that the resulting effective migration surface mirrors the outcomes of genetic differentiation detected by the *F_ST_* analyses (Figure 2); a higher effective migration was locally found in northern, northeastern and northwestern France among departments belonging to the same geographical areas, while a major barrier was discovered along the western side of France.

**Figure 2.**
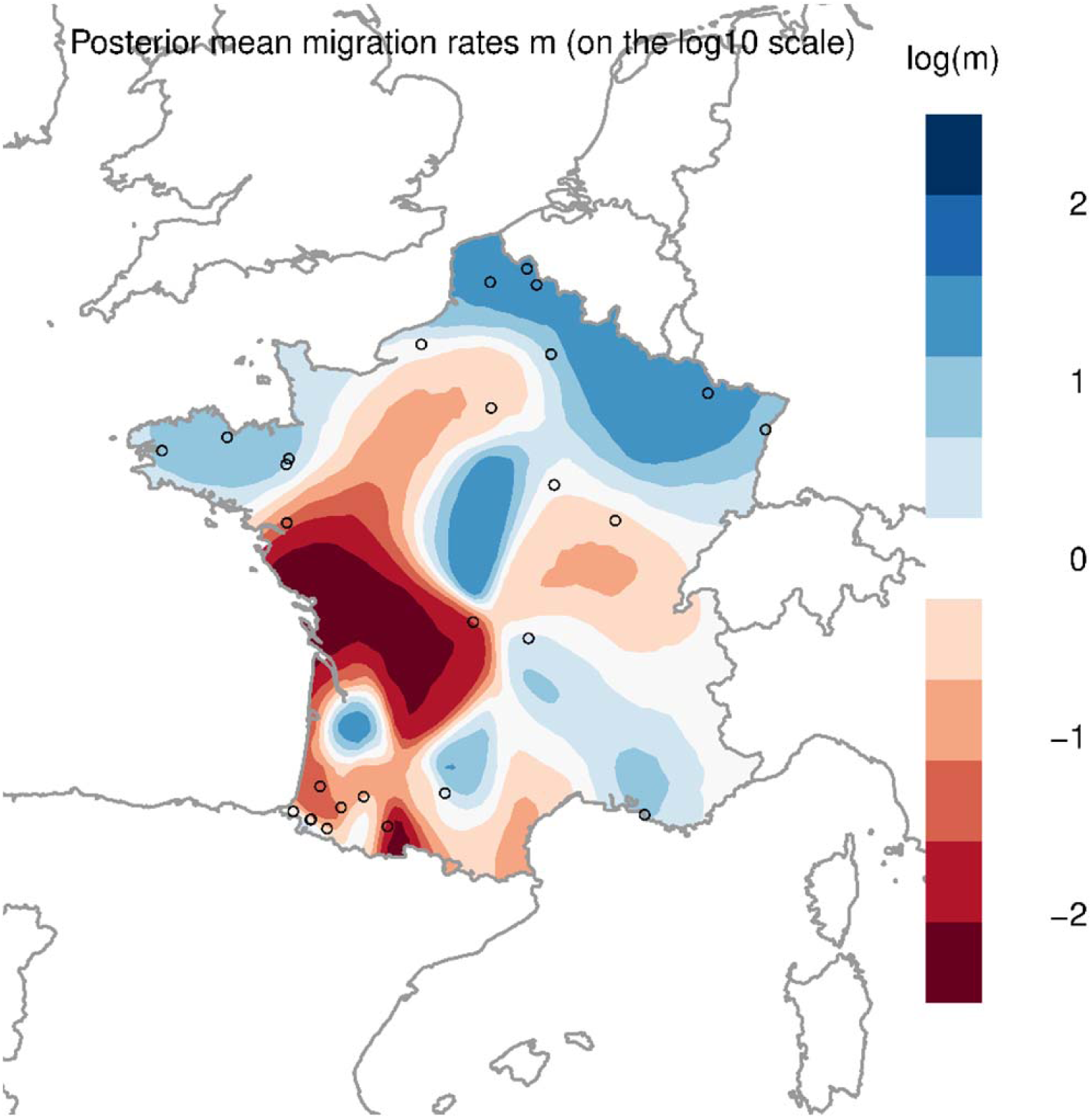
EEMS plot based on 395 French samples (Dataset A). Different shades of the same color represent differential levels of high (blue) or low (red) effective migration rates. The zero value indicates the average effective migration rate. Geographical locations for the different departments are averages of the coordinates among samples.

### Haplotype sharing patterns within France

Using haplotype-based methods (Dataset A), we looked for patterns of haplotype sharing, illustrating relations between departments. In this first step, cutting the fineSTRUCTURE tree at the very base, allowed us to describe a fine scale haplotype sharing distribution on a departmental scale; the outcome is a picture of the haplotype configuration within France (Figure 3). The resulting map shows finer-grained detail: we can define at least four distinct groups, plus a more widespread component. In the southwestern corner, the Basque samples clearly separate from the Gascon ones. In the northwestern vertex, the Breton departments exhibit their very own haplotypic signature, in agreement with the lower level of differentiation detected with the *F_ST_* analysis and the higher internal effective migration rate detected with EEMS. The same was found for the northern and northeastern departments that display a clearly shared haplotypic configuration. The southwestern department of Haute-Garonne (HG) and the southeastern one of Bouches-du-Rhône (BdR) present higher frequencies for some local haplotypes that in other departments reached only lower frequencies. Otherwise, a more generally spread French haplotypic background is found on the north-south axis.

**Figure 3.**
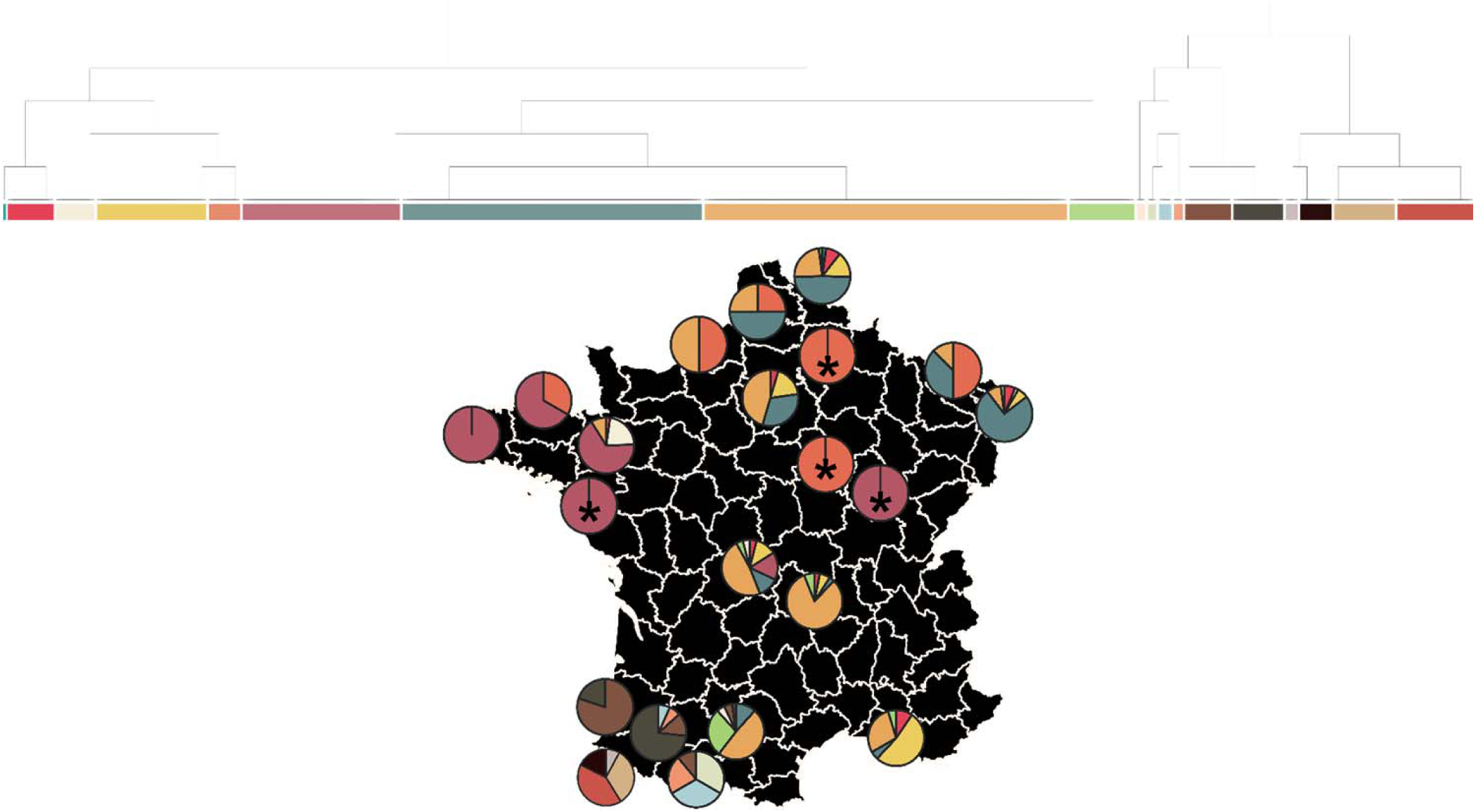
Pie charts showing the spatial distribution of haplotypes inferred by the fineSTRUCTURE tree. Each pie chart is a department, while colors correspond to the clusters described in the tree above the map. See Figure 1 for department names. Asterisks indicate departments with only one sample.

### Sources of gene flow into France

When we added external sources from the surrounding populations (yellow dots in Supplementary Figure 3) to describe allele-based genomic components with ADMIXTURE (Figure 4), the configuration observed pointed to a general homogeneous picture. The only exception was represented by the samples belonging to the Breton departments whose configuration was more alike to that in the Irish, Scottish, and English groups. Moving through the different K ancestral components, this behavior clearly characterizes the northwestern departments, separating them from the rest of the French groups since the very first K ancestral components (Figure 4). Thus, we formally tested for admixture events using the f3-statistics with the test groups being the different departments, and the external surrounding populations as sources. We only retained the negative f3 values for those departments represented at least by two individuals. Results are shown in Supplementary Table 2 were only significant Z-scores < −3 are reported, while results for those departments passing all the requested filters but with higher Z-score values are shown in Supplementary Table 3. Notably, in 9 departments, a combination of sources that was highly significant was Ireland-Southern Italy.

**Figure 4.**
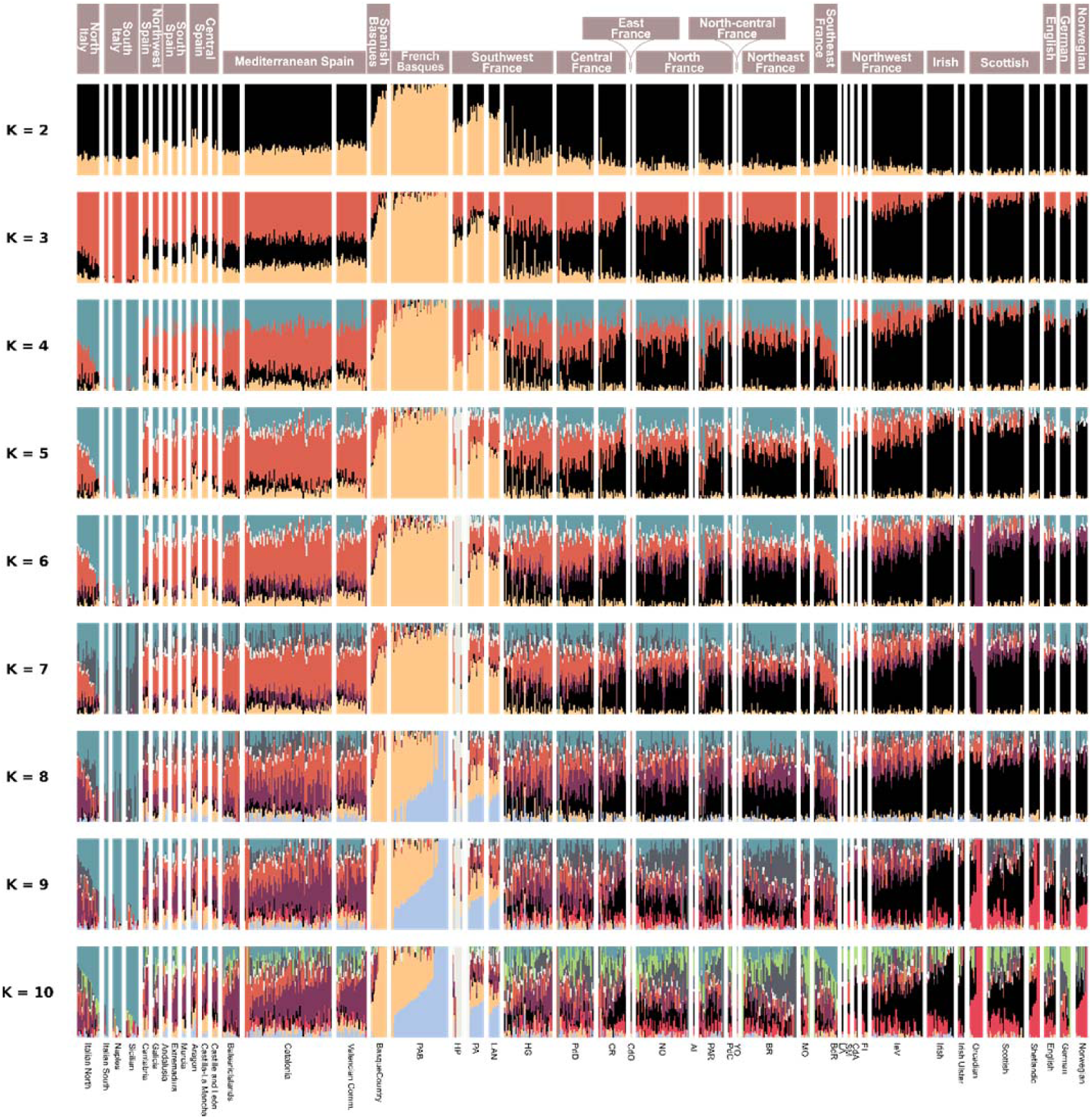
ADMIXTURE results from K=2 to K=10 for the 395 French samples (Dataset A) divided in nine major groups, and 12 groups representing external sources from surrounding countries; the lowest cross-validation error was found with K=2.

### Haplotype sharing patterns with external sources

Based on the haplotype sharing with external sources it was possible to redefine the French haplotype configuration. After merging the 395 French samples with the 1132 external ones (Dataset C), we first defined the external groups by silencing France when rerunning ChromoPainter and fineSTRUCTURE. The result was represented by 35 different external groups (Supplementary Figure 8a). Secondly, focusing on our target, we redefined the French internal clusters using the 35 external ones as “continents” when running fineSTRUCTURE (Supplementary Figure 8b). The 13 different clusters we found within France were then represented as separate maps (Figure 5); each map in the figure is a heatmap showing the number of samples falling in the different departments. Out of 13 groups, 10 satisfied the conditions of having at least 10 individuals and a major geographical area with a number of subjects corresponding to more than 50% of the entire cluster. These conditions allowed us to name each cluster based on the fact that a specific area was more represented than others in terms of sample size. The exclusion of three clusters did not impact the analysis, since only 8.35% of the French samples were then not included as target in the following analyses with GLOBETROTTER. As in the analysis described in the previous paragraph, even in this case France appeared to be organized in few major areas of interest. As shown in Figure 5, the Northwest presented two main groups (B1 and B2), the Southwest divided in Basque (Bas) and Gascon (G1 and G2) groups, the Northern (CN) and Northeastern (NE) areas, the Southeast (SE), and a central/southwestern part of France (CSW1 and SW). These ten main areas represented the targets for the GLOBETROTTER analysis that we used to describe the ancestry profiles, the admixture events, and their dates.

**Figure 5.**
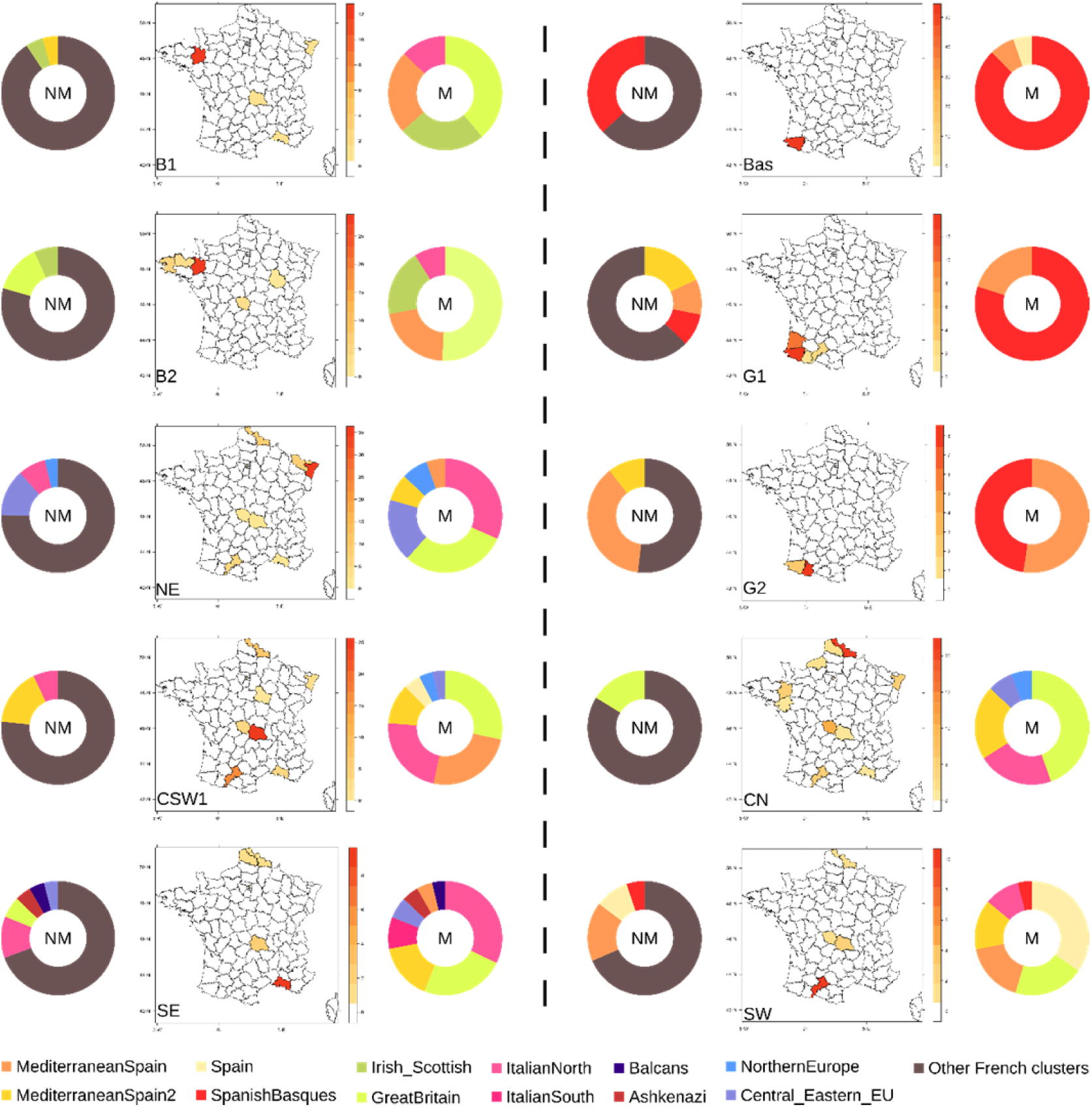
Ancestry profiles for 10 French targets. Each map is a target defining a specific major area of the French territory. On the left of each map, the donut chart is representing the ancestry profile for the not masked analysis (NM); on the right the same analysis has been masked (M). The different colors represent proportions of haplotype sharing with a specific source (only contributions above the 2.5% are shown); sources are defined in supplementary Figure 8. In the NM analysis, the brown color refers to contributions coming from other French groups (cumulative value). Target names stand for: B1 and B2, Brittany; NE, NorthEast; CSW1, Central-SouthWest; SE, SouthWest; Bas, Basques; G1 and G2, Gascons; CN, Central-North; SW, SouthWest.

### Ancestry profiles and dating admixture events

The results from the application of the nnls algorithm are displayed in Figure 5; on both sides of each target the ancestry profiles are represented as doughnut charts (on the left the results from the NM analysis, on the right the ones for the M one). The different colors represent proportions of haplotype sharing with specific sources (only contributions above 2.5% are shown). In the NM analysis, it is possible to appreciate how the haplotype sharing with other French sources (brown color) represents the highest proportion for all the different targets. When masking the French component, more refined patterns of contributions from external sources are detected. With the only exception of the southwestern targets (G1, G2, and Bas), the remaining ones show a higher contribution from north Italy and Great Britain. Apart from these common signal, it is possible to highlight contributions from those neighboring populations that are more geographically close to specific areas within the French territory. The southwestern targets (G1, G2, and Bas) received more from the Spanish side, the northwestern targets (B1 and B2) share more with the external cluster source named *Irish_Scottish* (with a proportion of 23.91% and 18.32% for the B1 and B2 targets respectively), the northeastern target (NE) is more connected to the external cluster source representing central and eastern European countries (receiving 17.64% from the source we named *Central_Eastern_EU*), as also from the *NorthernEurope* cluster source (which contributes 7.35% and 5.78% to the NE and CN targets, respectively). The southeastern target (SE) is mostly connected to the Italian sources and other Mediterranean countries, and the central/southwestern target (CSW1) clearly received more from both Spain and Italy.

As explained in Supplementary note 1, GLOBETROTTER provided evidence of admixture for 8 out of 10 targets, and for 5 of them we could also describe the dates and the sources of admixture as shown in Supplementary Figure 9. For three targets GLOBETROTTER gave *one-date* as result, while for the remaining two *one-date-multiway* was detected. In each case, only one date of admixture was detected; for the *one-date* groups a single admixing couple of sources was described, while two couples of sources were presented in the case of *one-date-multiway*. For a better interpretation of the results, consider the caption from Supplementary Figure 9.

### Relations with ancient populations

In the analysis with Dataset D, we first explored the position of France in the context of other modern populations, and then we focused on the relation with a set of ancient samples from different periods. In Supplementary Figure 10, panel A shows the PCA with the modern samples; France (white circles) is located in a position that mirrors its geographical situation, in between British, Irish, Mediterranean, central and eastern European samples. In panel B, a set of ancient samples was projected into the modern genetic space. In this second PCA, most of the French individuals are close to the Steppe and the Late Neolithic Bronze Age (LNBA) European samples, with some subjects connecting with the Anatolian Neolithic and the Early Neolithic Eurpean groups, and few others with the Europe Middle Neolithic and Chalcolithic (Europe_MNChL) samples. Results from the ADMIXTURE analysis are reported for the lowest cross-validation error detected (K=4 in Supplementary Figure 11). At this level, four ancestral components are clearly visible: the hunter-gatherer (HG) ancestry (principally represented by the Scandinavian HG, in pink), Neolithic (mostly Anatolian and then European, in green), the Iran Neolithic (black), and Natufian (purple). Again, the proportion of these components in France is intermediate between those in Southern and Central European groups. It is especially the Natufian component that seems to act as a discriminant factor, not only inside France where it is virtually absent with few exceptions on the Mediterranean side, but mostly among the various modern groups. Outgroup f3-statistics in the form of f3(Ancient, X; Mbuti) allowed us to quantify for each X modern group the amount of shared drift with different ancient populations. Figure 6 shows the outcome for these statistics, with a focus on the shared drift with the three main European ancestral components: Western, Eastern, and Scandinavian hunter-gatherers, European Neolithic farmers, and the European Bronze Age steppe component. In most cases, French populations fit the expected pattern of distribution in the wider panorama of the European area. However, the European Neolithic component seems to be higher in the SW of France, while Brittany carries a proportion of HG ancestry that is higher than elsewhere in France but closer to the values in the British Isles.

**Figure 6.**
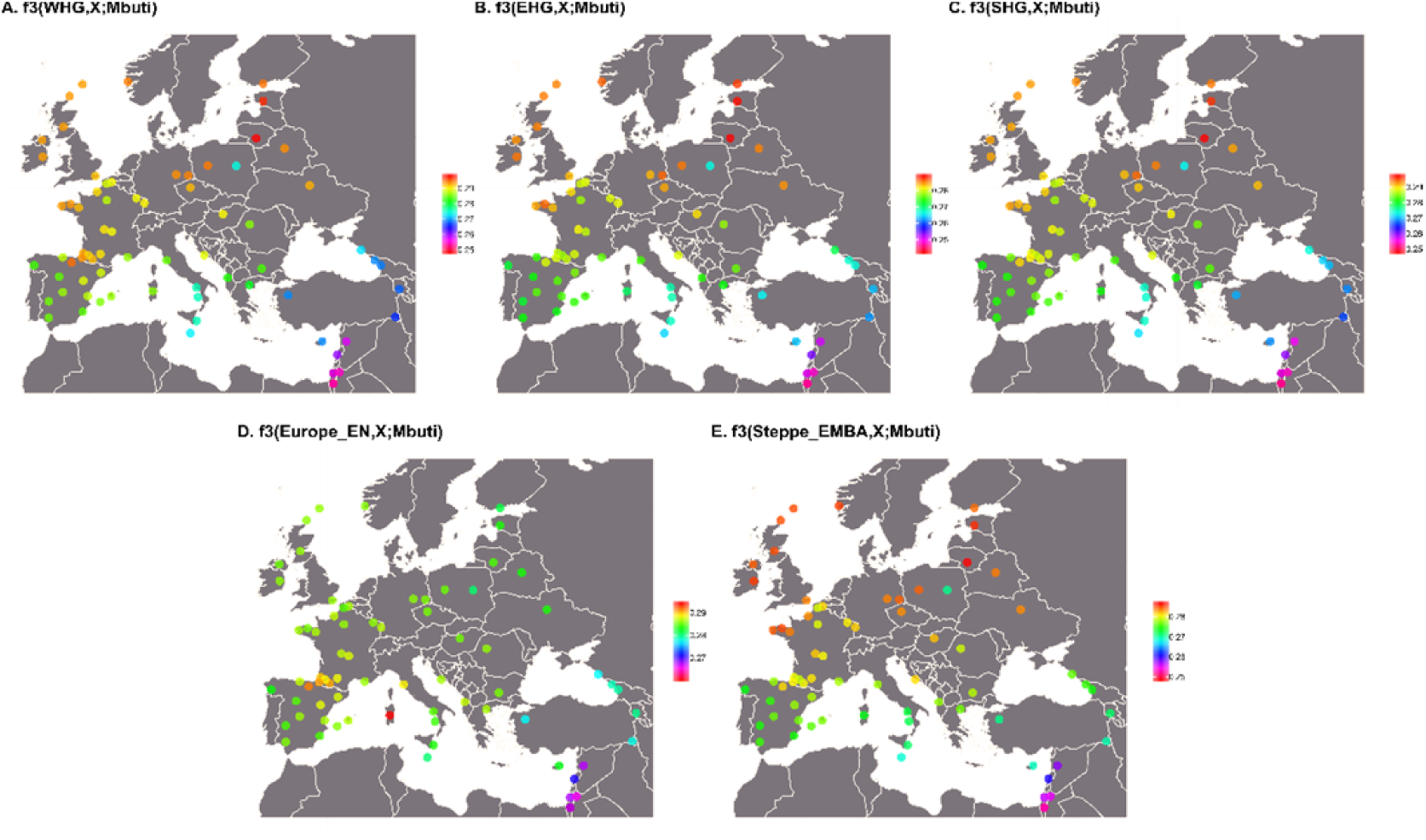
Maps showing the distribution of the shared drift between different ancestral populations and the modern ones (X in the f3 statistics). Panels: **A)** f3(Western Hunter Gatherers,X;Mbuti), **B)** f3(Eastern Hunter Gatherers,X;Mbuti), **C)** f3(Scandinavian Hunter Gatherers,X;Mbuti), **D)** f3(Europe_Early Neolithic, X;Mbuti), **E)** f3(Steppe Early Middle Bronze Age,X;Mbuti). In France, departments with less than two individuals are not shown.

## Discussion

We have used both allele frequency and haplotype-based methods in order to describe the internal structure of pre-20th century Metropolitan France. While the first yielded a more homogeneous landscape, the latter unveiled patterns of local differentiation with some connections with the surrounding European populations. Furthermore, we explored patterns of genetic continuity with ancestral populations, contextualizing France in the wider European panorama. In previous works about France, samples were differently arranged into the geographical space and no consensus had been reached on what subdivision was more appropriate; apart from the peculiar military districts ^3^, historical provinces ^4,6^ and old regions ^8^ are the most used so far. Thus, our first goal was to search for the best geographical level of genetic stratification before arranging our samples on a map. After the French Revolution in 1790, in order to weaken the old loyalties, the ancient provinces of France were subdivided into departments, whose overall configuration has been mostly conserved so far ^30^. Furthermore, in 1982, a system of 22 regions was established by grouping different departments into wider areas ^31^. However, in 2016, the number of the regions was reduced to 13, with the consequent rearrangement of the departments ^32^. Given this background, our AMOVA results provide evidence that regions, as a new internal reorganization, are not a suitable model for the genetic compartmentalization and point to the absence of any contribution to the total genetic variation, possibly implying that regions are separating genetically similar departments into different groups. On the other hand, departments, as result of a more conserved internal geographical structure, represent the best minimal unit of genetic stratification.

### Dissecting the *Hexagone*

Principal component analysis on allele frequencies revealed the expected Basque differentiation, adding Gascons in SW France as a population closely related to them, while the rest of France appeared relatively homogeneous. However, EEMS results pointed to the existence of other barriers to gene flow, particularly between NW France (Brittany) and the rest, while other areas acted as corridors, in central France and along the N and NE borders (Figure 2). It should not be excluded, though, that unsampled regions caused some possible artifacts ^33^. It was with fineSTRUCTURE that we could really define a fine scale internal subdivision of France (Figure 3). A general widespread French haplotypic background moving through the north-south axis was detected; possibly the overall homogeneity found with the principal component analysis can be linked to the fact that, on an allele frequency scale, such widespread pattern may represent a confounding factor. Indeed, only the two Southwestern groups (Basques and Gascons) were not reached by this common French haplotypic background. Particular haplotype sharing patterns could also be observed along the north and northeast of France, in the southeast, and among the northwestern departments.

In order to understand whether these internal patterns of differentiation are due to recent events or whether they reflect a more ancient history, we relied on different analyses obtaining distinct information. On the one hand, we looked at the relation with modern external populations, exploring both allele-frequency (ADMIXTURE and f3-statistics) and haplotype-based methods (using GLOBETROTTER, we described the ancestry profiles for 10 different French targets, defined by the haplotype sharing with external sources, and provided a date of admixture events for 5 of them). On the other hand, we looked for the continuity between modern France and ancestral populations from different times.

### France, *carrefour* of Europe

An ADMIXTURE plot (Figure 4), and a PCA with reference populations (Supplementary Figure 10A) place most French populations as similar to their geographic neighbours, namely the British Isles, Central Europe, Spain and Italy, in accordance with the general observation in Europe of geographic distance as the main predictor of genetic distance ^34,35^. This may explain an apparently surprising outcome of our work: 9 out of 22 distinct targets in f3 statistics we tested against different external sources gave significant results with the lowest Z-scores detected for the same couple represented by the South Italian and Irish sources. Z-scores lower than -3 indicate that our test populations are admixed from sources not necessarily identical but related to the sources we used in the analysis ^11^. Interestingly, these results found support in the outcome from the ancestry profiles we carried out with the Dataset C. The ancestry profiles described in Figure 5 are informative of differential migratory patterns ^36^ into each of the ten French genetic targets. The ancestry profiles are a way to describe the genome of each one of the ten French target as a mixture of the genomes from other groups, without inferring any particular admixture event ^37^. With this analysis, each target is described as a composition of different proportions of haplotype sharing with other sources, excluding the contribution of the group that we want to explain (no self-copying allowed). Following the previous results from the f3-statistics, in the M analysis we found that 7 out of the 10 targets we tested were mostly described by high proportions of haplotype sharing with both Italy and the British Isles. Furthermore, the NM analysis highlighted the presence of a very strong shared French component, possibly reflecting the result of a higher intermixing between individuals from the different parts of modern France.

An additional dimension to the central genetic position of France in Western Europe is given by the comparison with a time transect of ancient samples. The ADMIXTURE results for dataset D (Supplementary Figure 11), as well as the projected PCA (Supplementary Figure 10B) place France again as intermediate between Southern and Central Europe. However, this pattern is locally nuanced, as discussed below. Thus, it appears that France has been operating as a crossroads for human migration in Western Europe since, at least, the Early Neolithic.

### Basques and Gascons

These groups clearly differentiated from the rest of France both with allele frequency and with haplotype-based methods. It is interesting to notice that the presence of two distinct groups in the Southwestern region stressed the outcome of the isolation the Basque-speaking group experienced, splitting from their non-Basque-speaking neighbors from the very same department (PA and PAB groups). This finding is in agreement with their recognized distinct cultural entity ^38^ and their genetic outlier position in the European landscape ^39^, as also with the lower internal levels of differentiation we detected with the *F_ST_* analysis, and the low effective migration rates evidenced by EEMS, resulting in a barrier to migration in the southwestern corner of France.

The ancestry profile for French Basques (Figure 5) reflects an almost exclusive component from Spanish Basques, with some minor contribution from two other source clusters in the Iberian Peninsula. Quite often, Spanish populations are modelled as the result of a Basque background plus external admixture ^40^, so it is not surprising that haplotypes found in Basques are also present in Spain. French and Spanish Basques, as well as other populations in NE Iberia, share also an increase in shared drift with Early Neolithic ancient samples (Figure 5D). The Basque singularity has often been explained as due to the persistence of an ancient gene pool, as old as the Late Glacial ^41^, or as the Pre-Neolithic ^42^, or as the Neolithic ^43^ (as our results seems to suggest), but a recent analysis of a large number of ancient Iberian samples ^44^ points to a more recent divergence, probably in the Iron Age, of the Basque population.

Gascons have been shown to be intermediate between French Basques and other French populations by PCA (Figure 1), and to carry a sizeable proportion of Basque ancestry (Figure 5). This could be the result of the postulated contraction of the Basque-speaking lands since the late Antiquity. Place names may indicate that Basque or languages similar to it may have been spoken in Aquitaine (SW France) south of the Garonne river ^45^.

### The Celtic connection

As shown by EEMS (Figure 2), a barrier to gene flow delineates the northwestern corner of France, indicating the presence of another distinct group represented by the Breton departments. This group was firstly detected, on a coarser scale, with the removal of the Southwestern samples (Basques and Gascons) from the first PCA, and its outstanding position is in agreement with different studies on both uniparental and autosomal markers ^6–9^. However, based on the fineSTRUCTURE results, in our work we detected a stronger evidence of differentiation based on haplotypic data. ADMIXTURE showed a connection to the Irish samples (Figure 4), which is also indicated by the ancestry profiles of the B1 and B2 targets, which showed higher proportions for the *Irish_Scottish* cluster source (Figure 5). The GLOBETROTTER analysis for determining the admixture dates pointed to some interesting results (Supplementary Figure 9). B2, the largest Breton target, gave signals of admixture around 700 CE, in the time frame of the British Celtic migrations (from Cornwall and south-west Britain) into Gaulish Armorica (then renamed Brittany) from the 3rd to 9^th^ centuries CE, with a higher flow between the 5th and the 6th centuries CE ^46^.This completely agrees with previous findings ^7–9^. Historical migrations from Ireland to Brittany are well recorded since the 4th century CE ^47^, as well as the emigration of Irish people during the War of Ireland (1641-1651) into the present day departments of Finistère (FI) and Côte d’Armor (CdA), within which a higher integration of the Irish immigrants is proved by records of marriage, birth and death certificates ^7^. Furthermore, a Celtic root for the Breton language links the Breton departments to the Insular Celtic languages from the British Isles ^48^.

Still, the connection may be more ancient. In Figure 6, we explore the three main European ancestral components ^49^: the pre-Neolithic hunter-gatherers, the European Neolithic farmers, and the European Bronze Age steppe. Observing the shared drift with the three hunter-gatherer groups (panels A, B, and C), it is possible to notice how the northwestern departments are mirroring the values shown by the British Isles, the Central-Eastern countries, and Northern Europe. Brittany is thus showing a signal of continuity with the British Isles which could be ascribed to a period older than the later Celtic migration. Always Brittany is acting as an outlier in the case of the shared drift with the Steppe Early and Middle Bronze Age group. In Figure 6 (panel E) it is possible to see how Brittany breaks the northeast-to-southwest decreasing gradient of shared drift. Even in this context, Brittany shows a continuity with the British Isles. Actually, this is consistent with the archaeological records and the development of a late Megalithic culture that characterized Ireland, Britain and Brittany in a period when other parts of Europe were experiencing the advent of metallurgy ^50^.

### Borderlands

The northeastern rim of France, and the Mediterranean southeastern region represent areas in the perimeter of the *Hexagone* that may have received particular genetic influences. In the ancestry profiles (Figure 5), the NE and SE targets exhibit the most complex genetic make-ups, with a diverse array of sources. The *Central_Eastern_EU* cluster source is mostly represented in the NE target, which includes the departments of Bas-Rhin and Moselle; this area recalls the long history of the Alsace-Lorraine territory: a fuzzy border between France and Germany for a long time, and only recently retroceded to France in 1945 ^51^.

The SE target (most abundant in the Bouches-du-Rhône department) copied from several Mediterranean sources (thus representing the target with more complexity). This area has been a corridor and a landing place for different Mediterranean peoples, since 600 BCE when Greeks established a colony on the Mediterranean coastline of France in the city of Massalia (present-day Marseille) ^52^. However, this Mediterranean connection may be older, since the late Epipaelolithic Natufian component (Supplementary Figure 11), which is found almost exclusively in the Mediterranean populations, is found in France in the highest frequency in the Bouches-du-Rhône department.

### Conclusions

In conclusion, according to our results, France is a genetic intermediate between Central, Eastern, and Northern Europe, with some influences from the Mediterranean countries on the southeastern coast. Analyses with both modern and ancient groups pointed to a clear separation of the southwestern groups (Basques and Gascons) and of Brittany from the rest of the French areas. The application of haplotype-based methods allowed us to look beyond the more homogeneous French haplotypic background, discovering connections with the neighbouring populations (e.g., French northeastern departments with central and eastern Europe), while analyses with ancestral populations strengthened the historical connection between Brittany and the British Isles.

## DATA AVAILABILITY

The genotypes of the samples typed for this manuscript can be downloaded from https://figshare.com/articles/France_Dataset/10008689 and https://figshare.com/articles/Naples_Dataset/10008731

## ACKNOWLEDGMENTS

We thank Inés Quintela and the National Genotyping Center (CEGEN – USC) for their assistance in genotyping the samples. Of course, this work would not have been possible without the kind collaboration of all the sample donors. Funding was provided by the Agencia Estatal de Investigación and Fondo Europeo de Desarollo Regional (FEDER) (grant CGL2016-75389-P), Agència de Gestió d’Ajuts Universitaris i de la Recerca (Generalitat de Catalunya) grant 2014 SGR 866, and “Unidad de Excelencia María de Maeztu”, funded by the MINECO (ref: MDM-2014-0370). SAB was supported by the Agencia Estatal de Investigación FPI grant BES-2014-069224.

## AUTHOR CONTRIBUTIONS

SAB and FC designed the study; SAB carried out the analyses and interpretations, which were discussed with FC and DC; ERL and DC provided samples and unpublished genotypes. SAB wrote a first draft of the manuscript, with contributions from FC and DC. All authors read and approved the last version of the manuscript.

## Supplementary Data

**Supplementary Figure 1.**
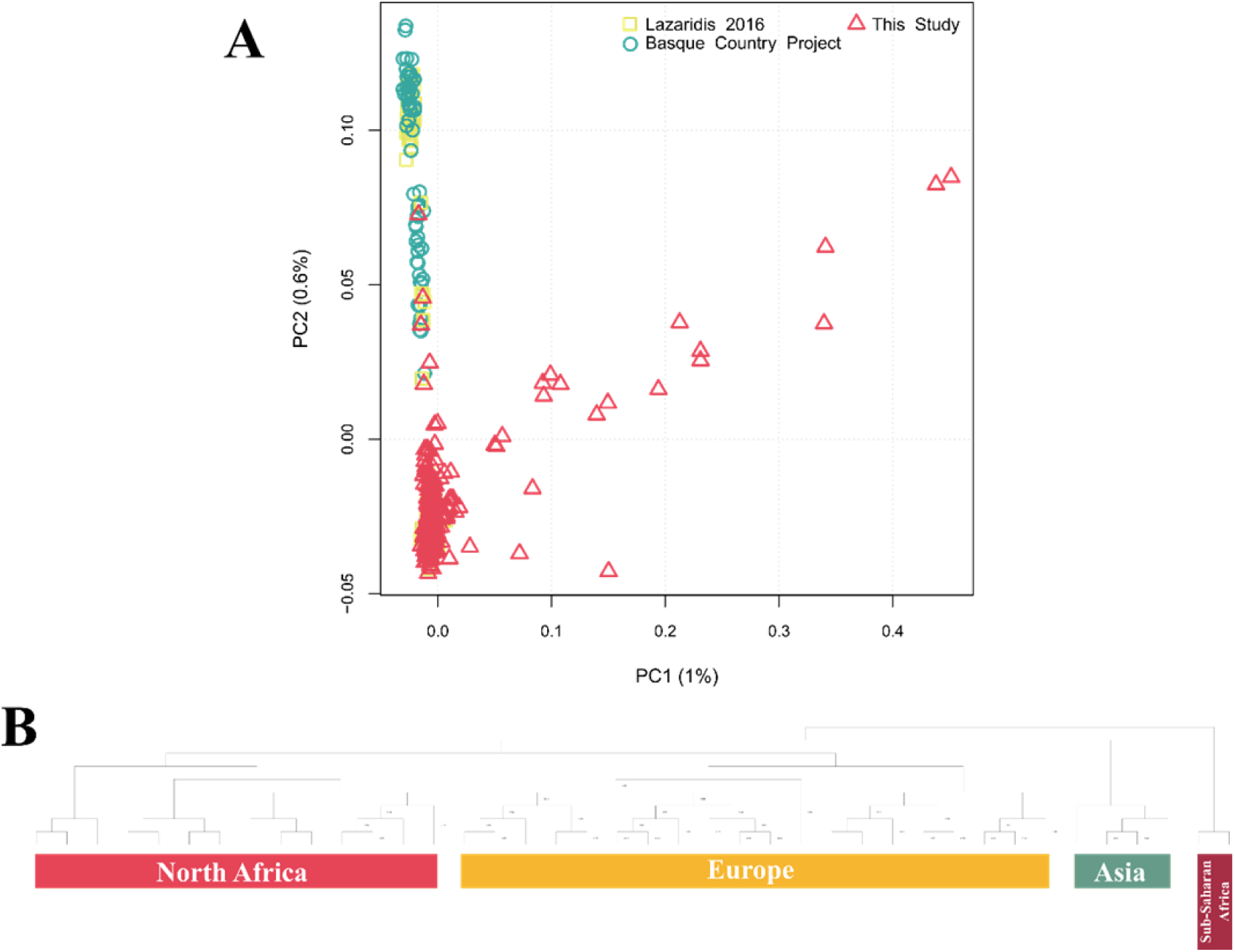
PCA with 415 French samples highlighted the presence of outliers clearly skewing the global distribution of the samples (**A**). We assessed the origin of those samples using ChromoPainter and fineSTRUCTURE in the context of external references from three worldwide populations (CEU, YRI, CHB) from the 1000 genomes project ^53^ and North African samples from published data ^11^. Four clusters were defined (**B**), assigning the majority of our samples (395) to the European cluster. The remaining 20 were outliers mainly belonging to the North African cluster (16 samples), 2 samples each were instead assigned to the Asian and the Sub-Saharan African clusters.

**Supplementary Figure 2.**
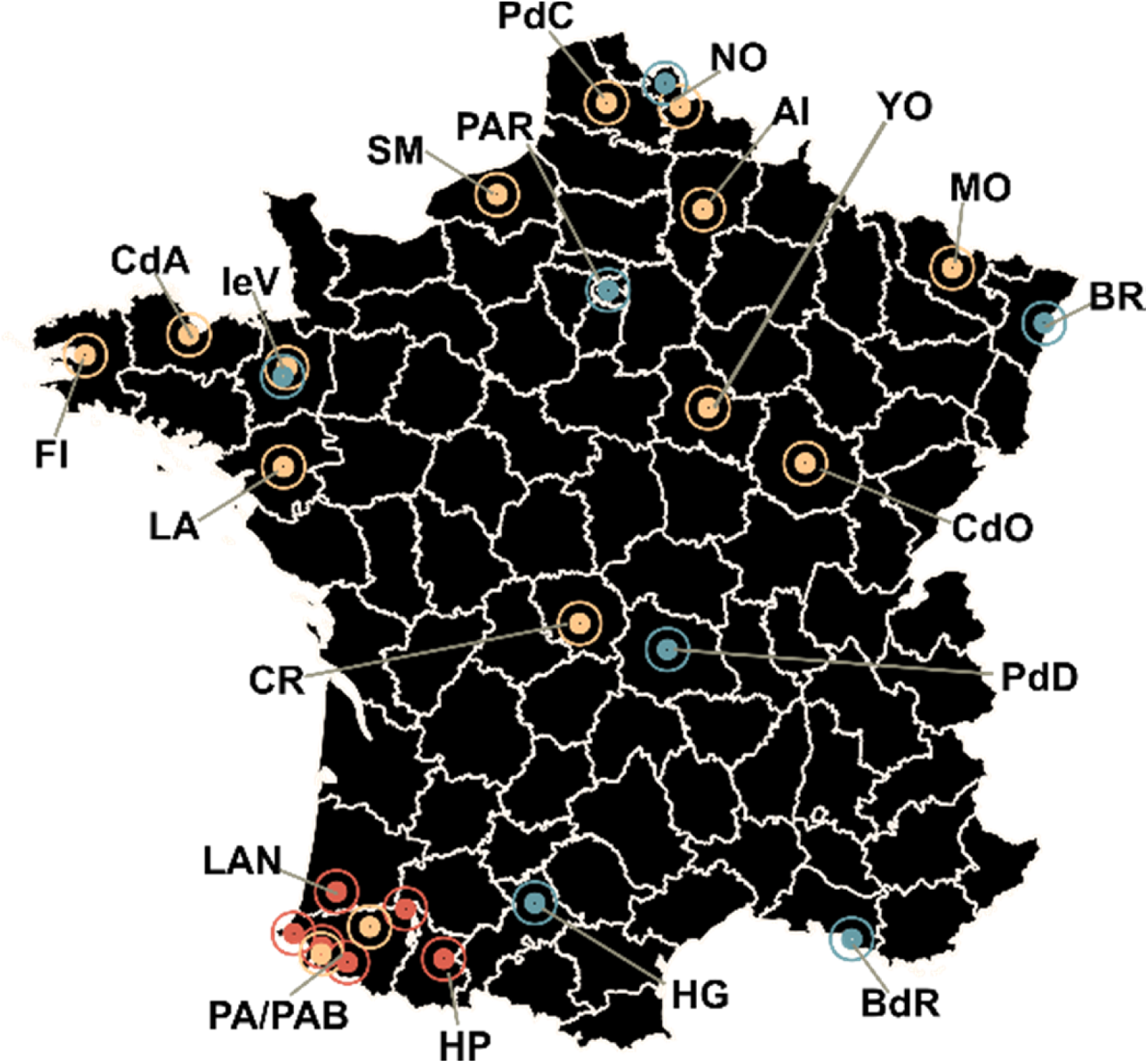
Map showing sample distribution among the different departments. Geographical coordinates are averages among samples. Different colors define the three datasets used in this work (blue dots correspond to the 256 samples genotyped for this work; yellow dots correspond to the 79 samples from Lazaridis et al., 2016; red dots correspond to the 60 samples from unpublished data). Sample size and acronyms for the departments are: PdD, Puy-de-Dôme (33); CR, Creuse (25); CdO, Côte-d’Or (1); AI, Aisne (1); NO, Nord (47); PdC, Pas-de-Calais (4); PAR, Paris (22); YO, Yonne (1); MO, Moselle (8); BR, Bas-Rhin (48); IeV, Ille-et-Vilaine (45); CdA, Côtes-d’Armor (3); FI, Finistère (5); SM, Seine-Maritime (2); LA, Loire-Atlantique (1); BdR, Bouches-du-Rhône (21); LAN, Landes (10); HG, Haute-Garonne (43); PA, Pyrénées-Atlantiques (15); PAB, Pyrénées-Atlantiques Basque (31); HP, Hautes-Pyrénées (*2*9).

**Supplementary Figure 3.**
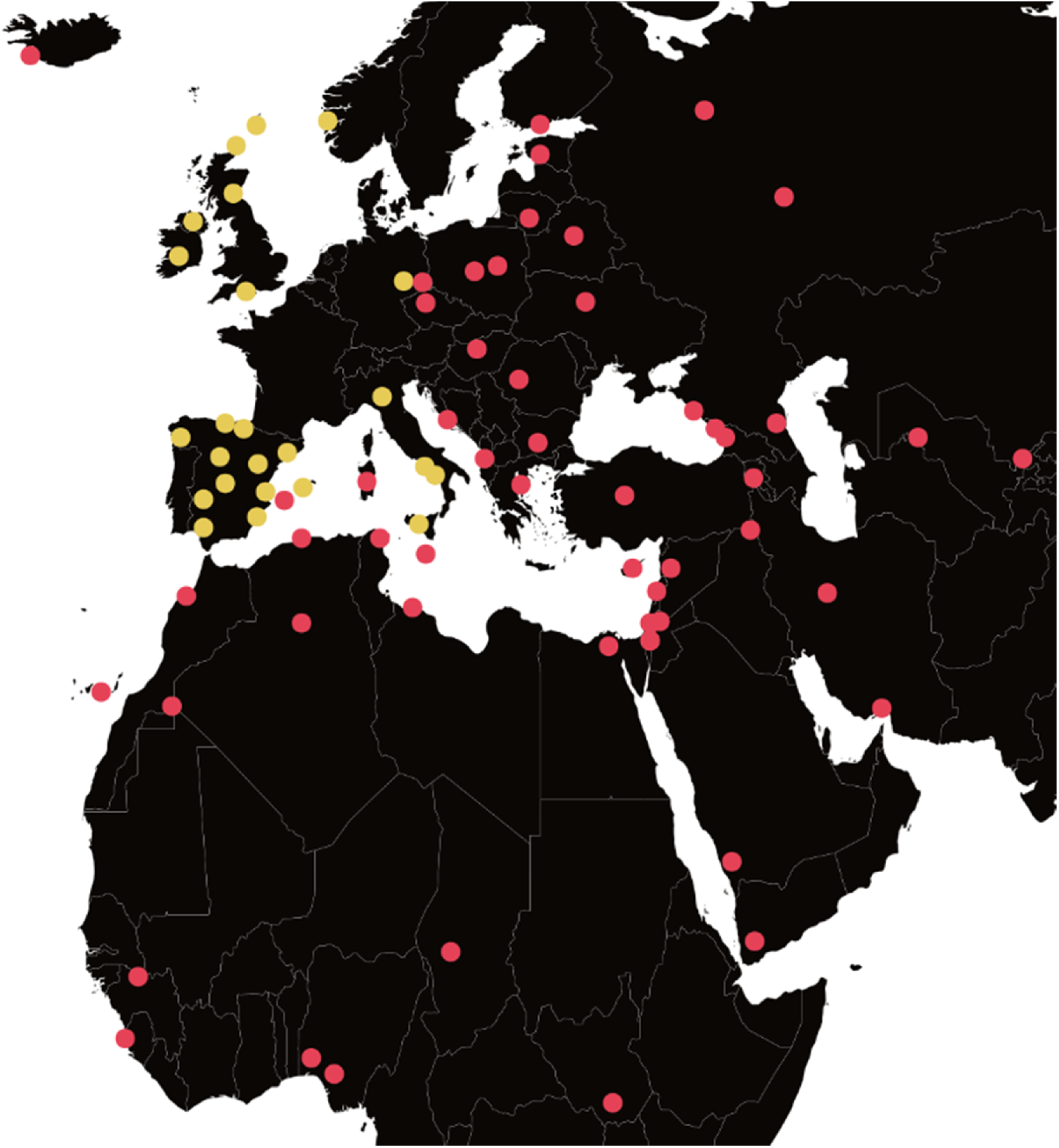
External group distribution. Average geolocation points for the 79 external populations are displayed. Yellow dots refer to the 333 samples included in the allele frequency analyses. Yellow and red points together represent the 1132 samples used in the haplotype-based analyses

**Supplementary Figure 4.**
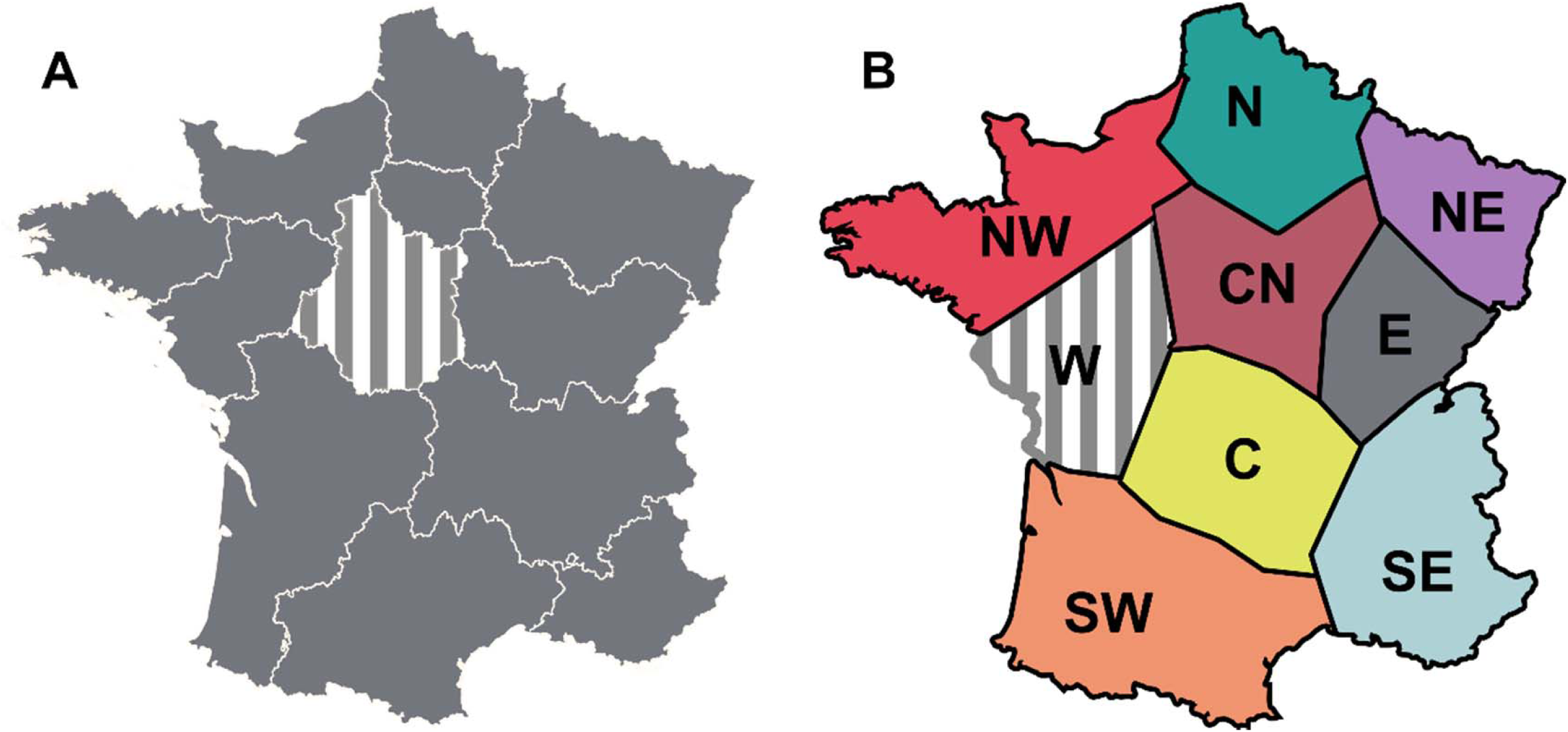
Higher hierarchical levels used in the AMOVA analysis for **A)** Regions and **B)** Areas. Grey vertical lines highlight unsampled zones. Acronyms for the Areas are: NW, Northwest; N, North; NE, Northeast; W, West; CN, Central North; E, East; C, Center; SW, Southwest; SE, Southeast.

**Supplementary Figure 5.**
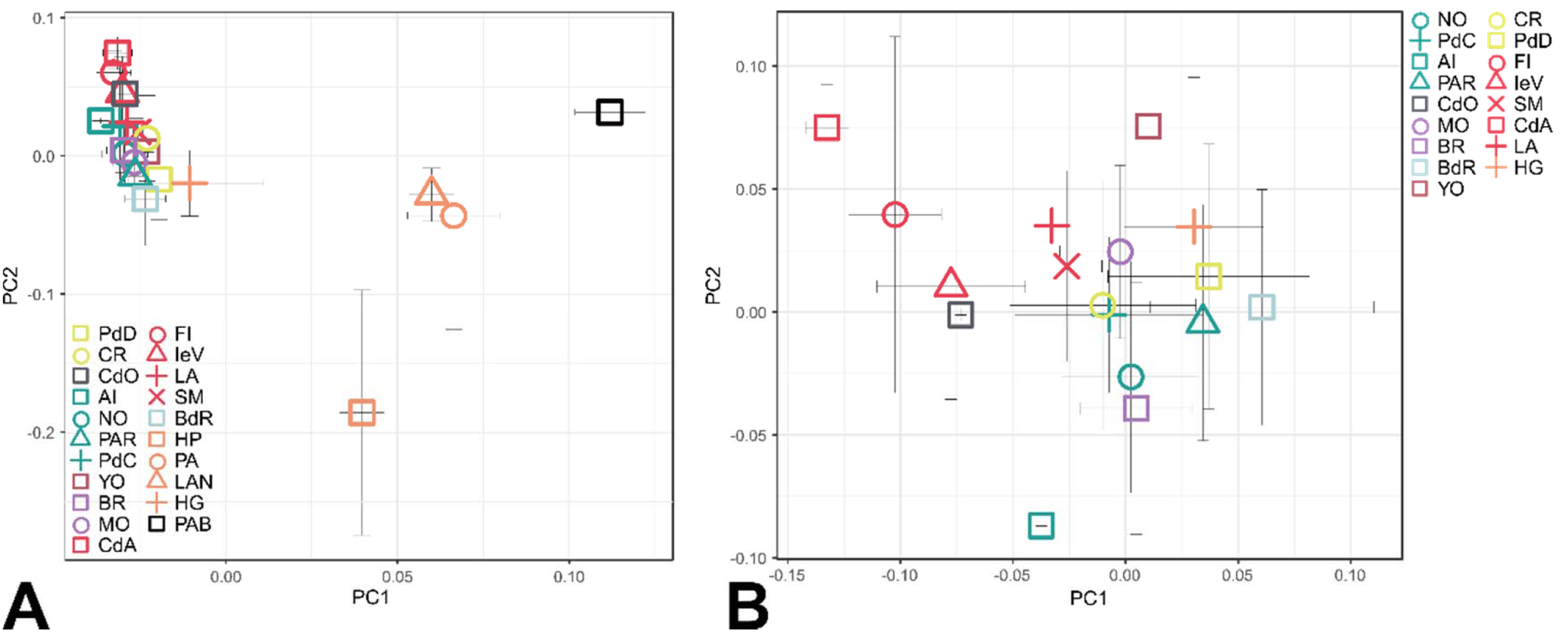
Averaged Principal Component Analysis with **A)** Basque and Gascon samples, and **B)** without them. Color and symbol codes are the same as in main Figure 2. For each group, each averaged eigenvalue is represented along with standard deviation bars for the two PCs.

**Supplementary Figure 6.**
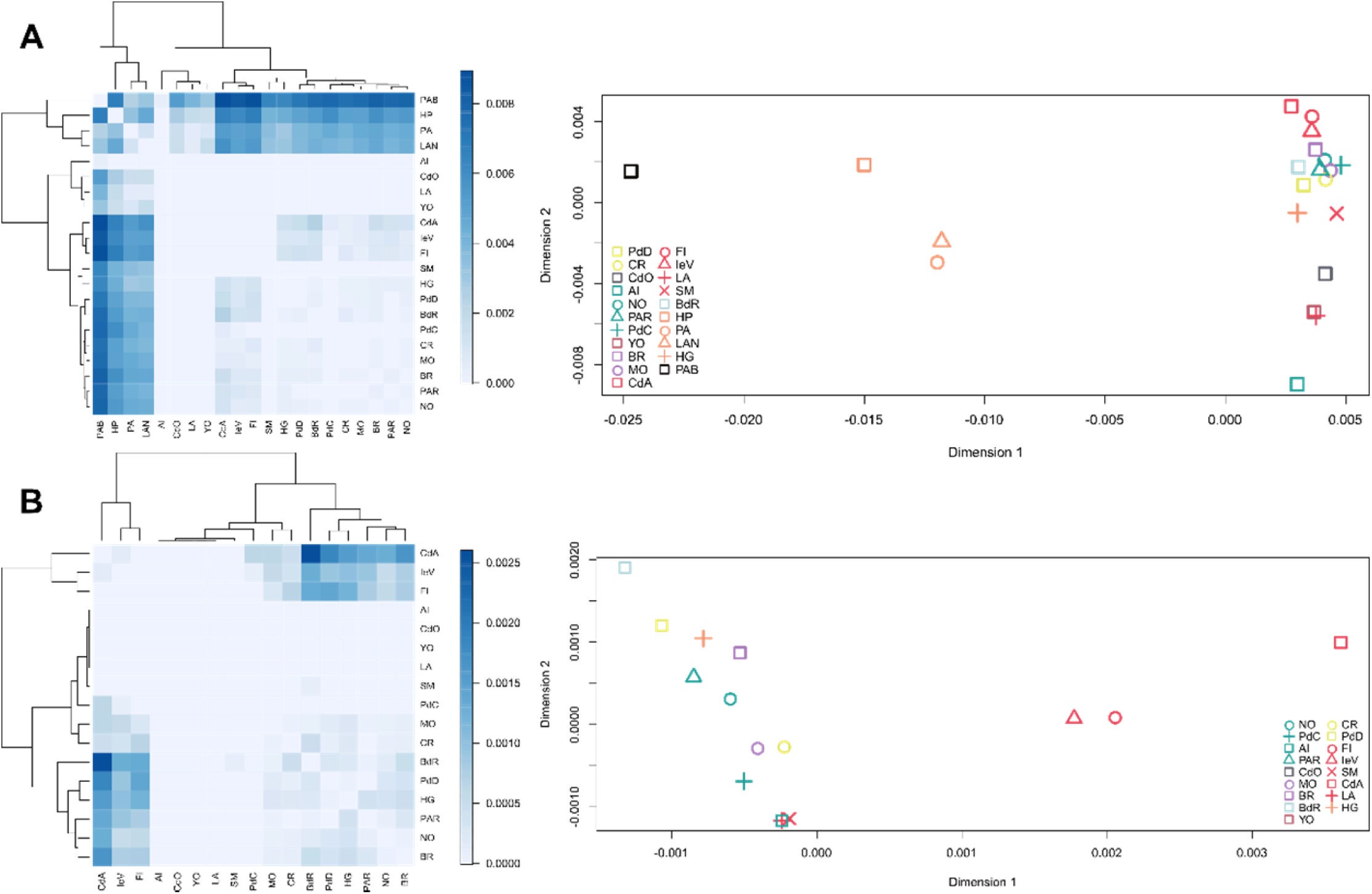
On the left: heatmap and dendrogram based on *F_ST_* matrices **A)** with the Basque and Gascon samples and **B)** without them. On the right: Multidimensional scaling (MDS) based on *F_ST_* values **A)** with the Franco-Cantabrian samples and **B)** without them.

**Supplementary Figure 7.**
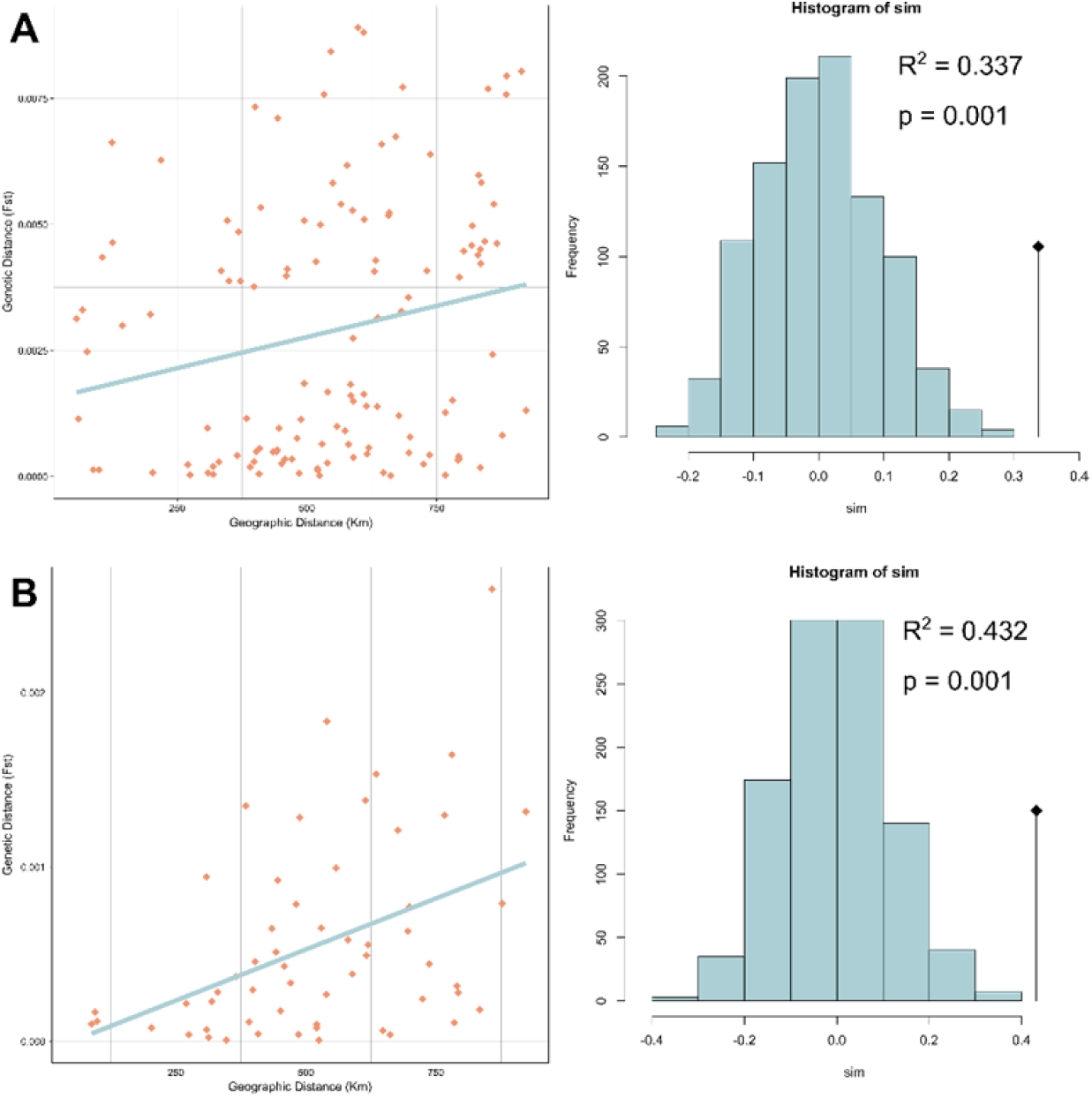
Mantel test of isolation by distance between the genetic (*F_ST_*) and geographic (in Km) distances **A)** with the Basque and Gascon samples and **B)** without them. R^2^ scores and p-values are within each figure.

**Supplementary Figure 8.**
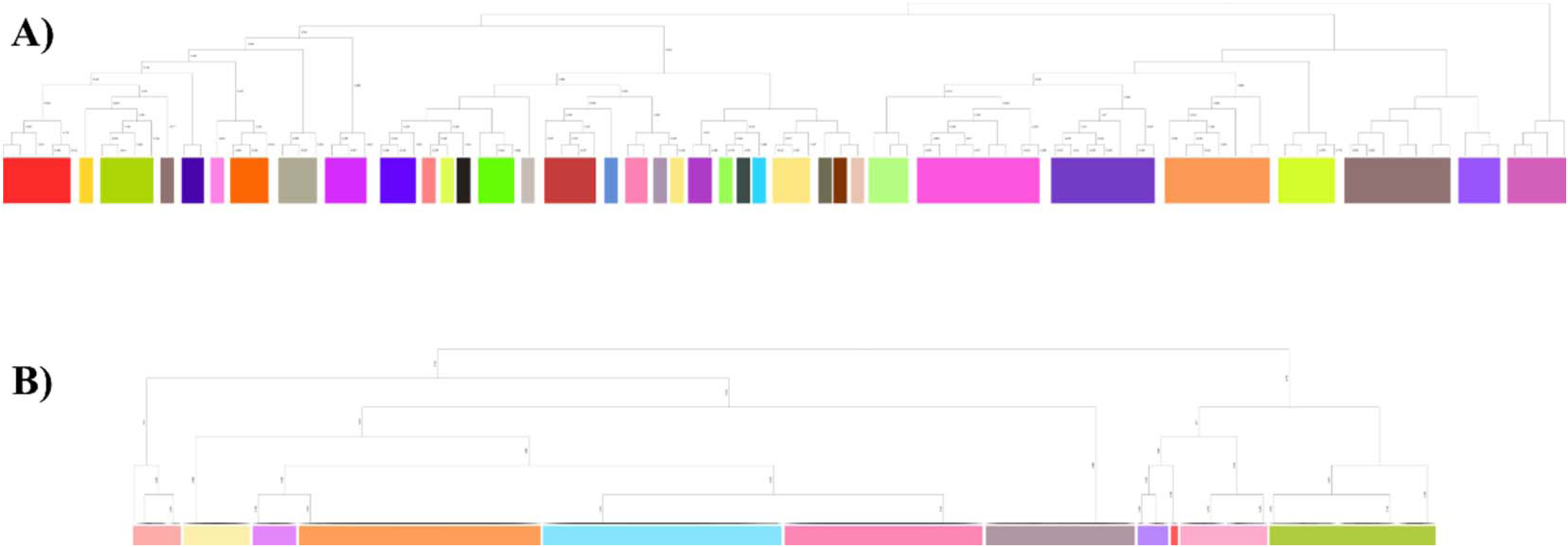
**A)** 35 clusters detected for the external samples when France was silenced in the rerun of CromoPainter and fineSTRUCTURE. **B)** 11 clusters detected within France using the “force file” option (-F) in fineSTRUCTURE.

**Supplementary Figure 9.**
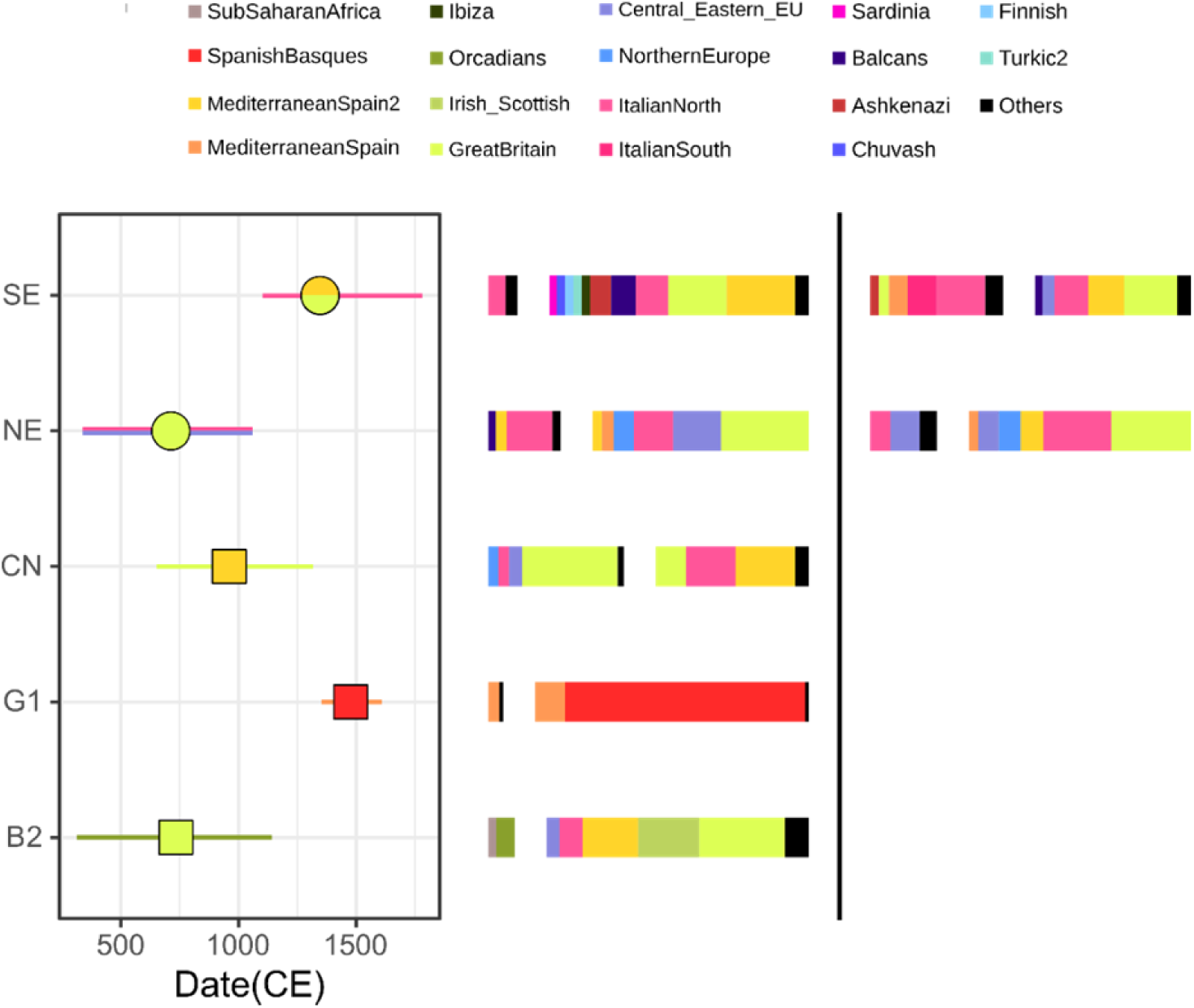
Dating results for 5 French targets according to the M analysis in GLOBETROTTER. In the left panel, squares refer to one-date, circles to one-date-multiway. The internal color refers to the highest surrogate’s value of the major source, while the color of the CI bars corresponds to the highest surrogate’s value of the minor source. Sources are represented as horizontal bars on the right side and are separated by a white space (together the sources account for the 100% of the values). In the one-date-multiway cases, two different sets of sources are presented and, where needed, both colors are represented for major and minor sources. Dates have been calculated as 1950-(g*N) where g=28 years and N is the calculated number of generations in the GLOBETROTTER analysis.

**Supplementary Figure 10.**
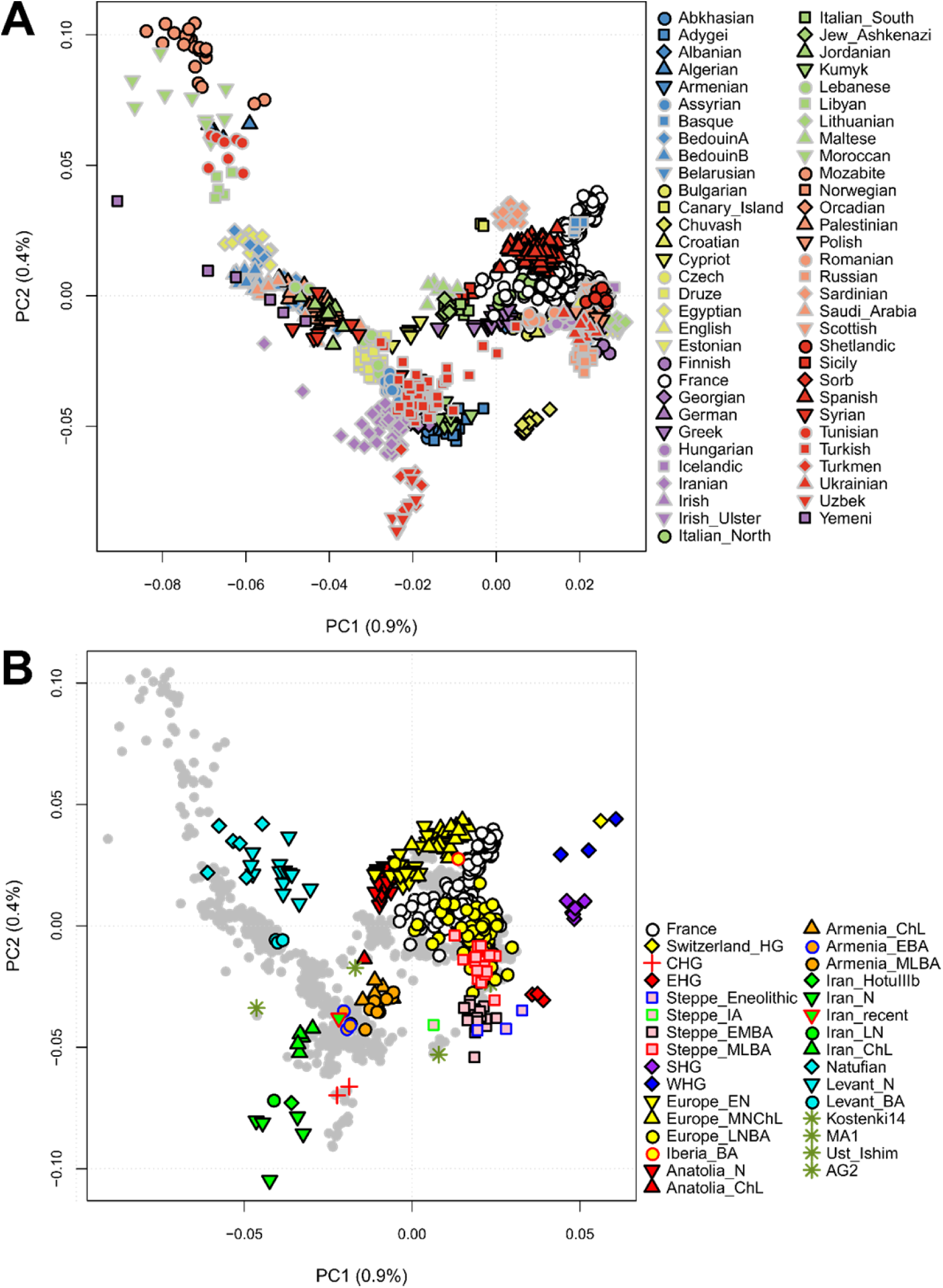
Principal component analysis with dataset D. **A)** Only modern samples; **B)** Projection of ancestral populations from different periods on top of the modern samples (grey dots; among the modern populations, only France is distinguishable as white circles).

**Supplementary Figure 11.**
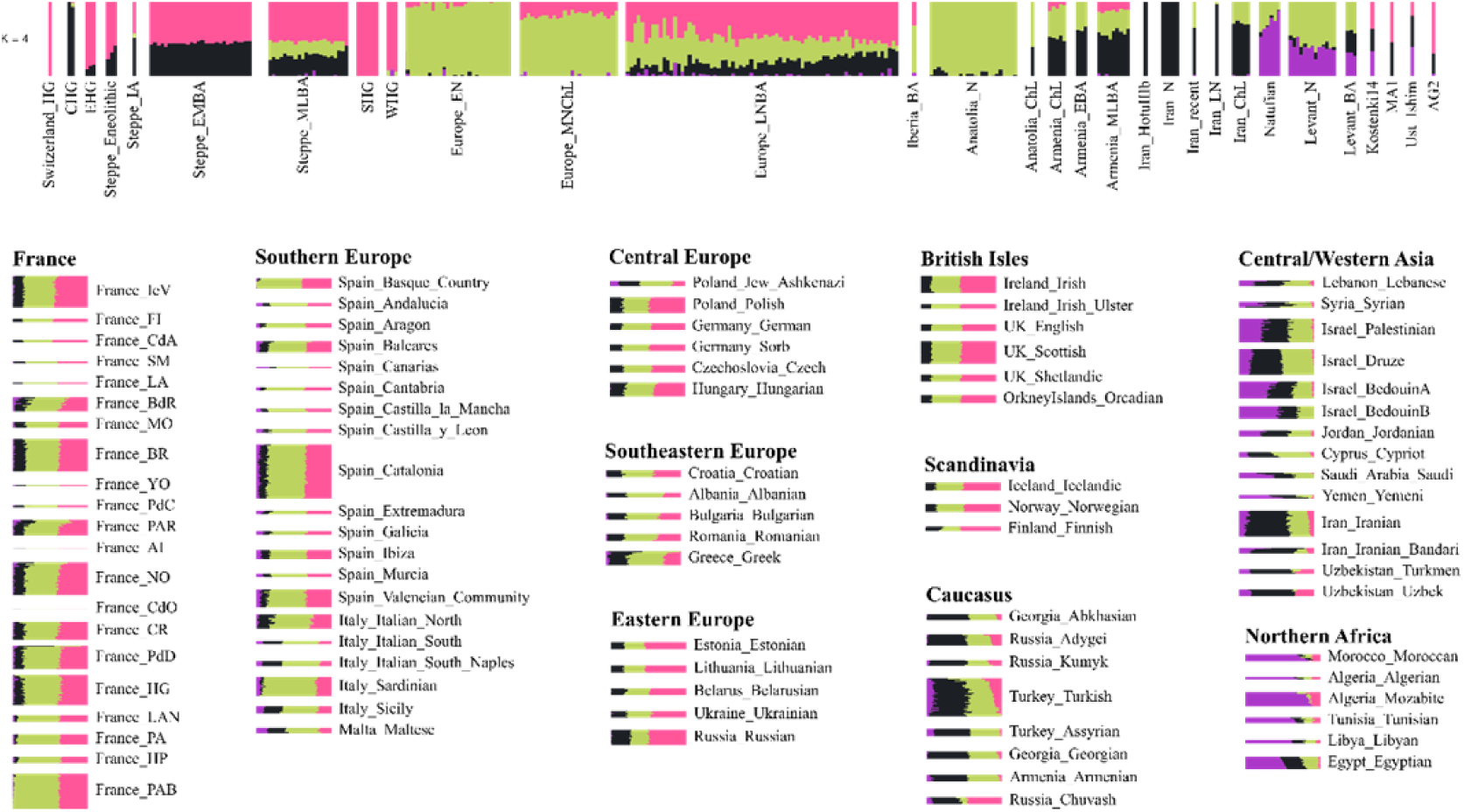
ADMIXTURE results for K=4 ancestral components using dataset D. Results for the ancient samples are on the top of the figure. Below, modern samples are organized according to major geographical groupings.

**Supplementary Table 1.**
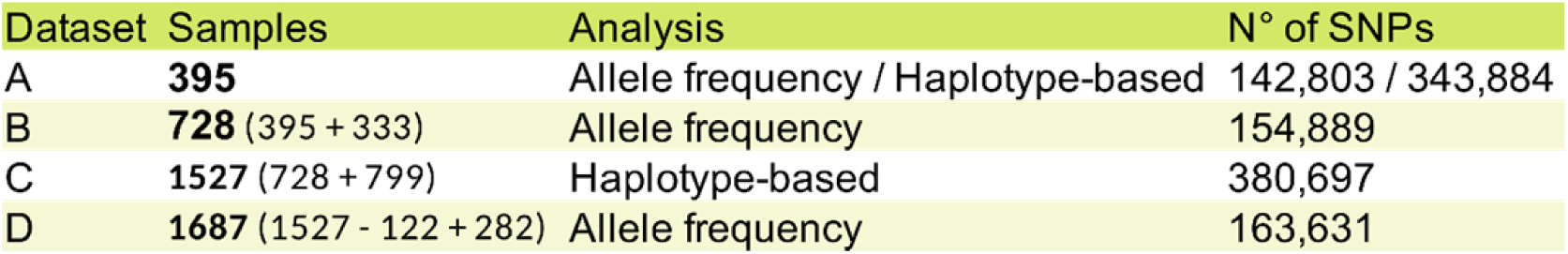
Summary of the dataset composition; both number of samples and number of variants are reported according to the analysis the dataset was used for.

| Dataset | Samples | Analysis | N° of SNPs |
| --- | --- | --- | --- |
| A | **395** | Allele frequency / Haplotype-based | 142,803 / 343,884 |
| B | **728** (395 + 333) | Allele frequency | 154,889 |
| C | **1527** (728 + 799) | Haplotype-based | 380,697 |
| D | **1687** (1527 - 122 + 282) | Allele frequency | 163,631 |

## References

1. Haine, W. S. The history of France. (Greenwood Press, 2000).

2. MacMaster, N. Colonial Migrants and Racism Algerians in France, 1900–62. (Palgrave Macmillan, 1997).

3. Kherumian, R., Moullec, J. & Nguyen, V. C. Groupes sanguins érythrocytaires A□, A□, BO, MN, Rh (CcDE) et sériques, Hp, Tf, Gm dans quatre régions militaires françaises. Bull. Mem. Soc. Anthropol. Paris 1, 377–384 (1967).

4. Cambon-Thomsen, A. & Ohayon, E. Practical Application of Population Genetics: The Genetic Survey “Provinces Françaises”. in Advances in Forensic Haemogenetics. Advances in Forensic Haemogenetics, vol 2. (ed. Mayr, W. R.) 535–553 (1988).

5. Cavalli-Sforza, L. L., Menozzi, P. & Piazza, A. The History and Geography of Human Genes. (Princeton University Press, 1994).

6. Richard, C. et al. An mtDNA perspective of French genetic variation. Ann. Hum. Biol. 34, 68–79 (2007).

7. Dubut, V. et al. mtDNA polymorphisms in five French groups: importance of regional sampling. Eur. J. Hum. Genet. 12, 293–300 (2004).

8. Ramos-Luis, E. et al. Y-chromosomal DNA analysis in French male lineages. Forensic Sci. Int. Genet. 9, 162–168 (2014).

9. Karakachoff, M. et al. Fine-scale human genetic structure in Western France. Eur. J. Hum. Genet. 23, 831–836 (2015).

10. Patterson, N. et al. Ancient Admixture in Human History. Genetics 192, 1065– 1093 (2012).

11. Lazaridis, I. et al. Genomic insights into the origin of farming in the ancient Near East. Nature 536, 419–424 (2016).

12. Martínez-Cruz, B. et al. Evidence of pre-Roman tribal genetic structure in Basques from uniparentally inherited markers. Mol. Biol. Evol. 29, 2211–22 (2012).

13. Biagini, S. A. et al. People from Ibiza: an unexpected isolate in the Western Mediterranean. Eur. J. Hum. Genet. 27, 941–951 (2019).

14. Purcell, S. et al. PLINK: a tool set for whole-genome association and population-based linkage analyses. Am. J. Hum. Genet. 81, 559–75 (2007).

15. Patterson, N., Price, A. L. & Reich, D. Population structure and eigenanalysis. PLoS Genet. 2, 2074–2093 (2006).

16. RStudio Team. RStudio Team (2015). RStudio: Integrated Development for R. (2015).

17. R Core Team. R: A Language and Environment for Statistical Computing. R Foundation for Statistical Computing, Vienna. (2018).

18. Dray, S. & Dufour, A.-B. The ade4 Package: Implementing the Duality Diagram for Ecologists. J. Stat. Softw. 22, (2007).

19. Wickham, H. ggplot2. (Springer New York, 2009). doi:10.1007/978-0-387-98141-3.

20. Wickham, H. Reshaping Data with the reshape Package. J. Stat. Softw. 21, (2007).

21. Kamvar, Z. N., Brooks, J. C. & Grünwald, N. J. Novel R tools for analysis of genome-wide population genetic data with emphasis on clonality. Front. Genet. 6, (2015).

22. Kamvar, Z. N., Tabima, J. F. & Grünwald, N. J. Poppr : an R package for genetic analysis of populations with clonal, partially clonal, and/or sexual reproduction. PeerJ 2, e281 (2014).

23. Alexander, D. H. & Novembre, J. Fast Model-Based Estimation of Ancestry in Unrelated Individuals. Genome Res. 19, 1655–1664 (2009).

24. Behr, A. A., Liu, K. Z., Liu-Fang, G., Nakka, P. & Ramachandran, S. pong: fast analysis and visualization of latent clusters in population genetic data. Bioinformatics 32, 2817–2823 (2016).

25. Petkova, D., Novembre, J. & Stephens, M. Visualizing spatial population structure with estimated effective migration surfaces. Nat. Genet. 48, 94–100 (2016).

26. O’Connell, J. et al. A General Approach for Haplotype Phasing across the Full Spectrum of Relatedness. PLoS Genet. 10, e1004234 (2014).

27. Delaneau, O. et al. Integrating sequence and array data to create an improved 1000 Genomes Project haplotype reference panel. Nat. Commun. 5, 1–9 (2014).

28. Lawson, D. J., Hellenthal, G., Myers, S. & Falush, D. Inference of Population Structure using Dense Haplotype Data. PLoS Genet. 8, e1002453 (2012).

29. Hellenthal, G. et al. A genetic atlas of human admixture history. Science (80-.). 343, 747–751 (2014).

30. Forstenzer, T. R. French Provincial Police and the Fall of the Second Republic: Social Fear and Counterrevolution. (Princeton University Press, 2016).

31. Sowerwine, C. France since 1870: Culture, Society and the Making of the Republic. (Palgrave Macmillan, 2009).

32. OECD. OECD Multi-level Governance Studies Multi-level Governance Reforms Overview of OECD Country Experiences. (2017).

33. House, G. L. & Hahn, M. W. Evaluating methods to visualize patterns of genetic differentiation on a landscape. Mol. Ecol. Resour. 18, 448–460 (2018).

34. Novembre, J. et al. Genes mirror geography within Europe. Nature 456, 98–101 (2008).

35. Lao, O. et al. Correlation between Genetic and Geographic Structure in Europe. Curr. Biol. 18, 1241–1248 (2008).

36. Leslie, S. et al. The fine-scale genetic structure of the British population. Nature 519, 309–314 (2015).

37. Hellenthal, G. Instruction Manual for “ GLOBETROTTER : a program for identifying, dating and describing admixture events in population data ” for GLOBETROTTER. 1–24 (2015).

38. Calafell, F. & Bertranpetit, J. Principal component analysis of gene frequencies and the origin of Basques. Am. J. Phys. Anthropol. 93, 201–215 (1994).

39. Rodríguez-Ezpeleta, N. et al. High-density SNP genotyping detects homogeneity of Spanish and French Basques, and confirms their genomic distinctiveness from other European populations. Hum. Genet. 128, 113–117 (2010).

40. Bycroft, C. et al. Patterns of genetic differentiation and the footprints of historical migrations in the Iberian Peninsula. Nat. Commun. 10, 551 (2019).

41. Pereira, L. et al. High-resolution mtDNA evidence for the late-glacial resettlement of Europe from an Iberian refugium. Genome Res. 15, 19–24 (2005).

42. Calafell, F. & Bertranpetit, J. A simulation of the genetic history of the Iberian Peninsula. Curr. Anthropol. 34, 735–745 (1993).

43. Günther, T. et al. Ancient genomes link early farmers from Atapuerca in Spain to modern-day Basques. Proc. Natl. Acad. Sci. U. S. A. 112, 11917–22 (2015).

44. Olalde, I. et al. The genomic history of the Iberian Peninsula over the past 8000 years. Science 363, 1230–1234 (2019).

45. Zuazo, K. El euskera y sus dialectos. (Alberdania, 2010).

46. Koch, J. Breton Migrations. in Celtic Culture : A Historical Encyclopedia 275–277 (ABC-CLIO, 2005).

47. Monnier, J. Chapitre 6 : L’immigration bretonne en Armorique. in Toute l’histoire de Bretagne (eds. Monnier, J. & Cassard, J.) 97–106 (Skol Vreizh, 1997).

48. Forster, P. & Toth, A. Toward a phylogenetic chronology of ancient Gaulish, Celtic, and Indo-European. Proc. Natl. Acad. Sci. 100, 9079–9084 (2003).

49. Lazaridis, I. et al. Ancient human genomes suggest three ancestral populations for present-day Europeans. Nature 513, 409–413 (2014).

50. Arias, P. The Origins of the Neolithic Along the Atlantic Coast of Continental Europe: A Survey. J. World Prehistory 13, 403–464 (1999).

51. Blumenthal, D. Alsace-Lorraine: A Study of the Relations of the Two Provinces to France and to Germany, and a Presentation of the Just Claims of Their People (Classic Reprint). (Forgotten Books, 2012).

52. Fine, J. V. A. The Ancient Greeks: A Critical History. (Belknap Press: An Imprint of Harvard University Press, 1985).

53. 1000 Genomes Project Consortium et al. A global reference for human genetic variation. Nature 526, 68–74 (2015).

54. Busby, G. B. J. et al. The Role of Recent Admixture in Forming the Contemporary West Eurasian Genomic Landscape. Curr. Biol. 25, 2518–2526 (2015).

