## Supplementary Note for "Reshaping the *Hexagone*: the genetic landscape of modern France"

**Supplementary note 1**

As previously pointed out [1], masking closely related groups allows to detect more refined admixture signals, and this is why we ran GLOBETROTTER twice: the first time we included also the French groups as surrogates (Not Masked analysis: NM), while the second time the French groups were excluded as surrogates (Masked analysis: M).

For the ancestry profiles, we used the options ‘prop.ind: 1’, ‘null.ind:0’, and ‘num.mixing.iterations:0’; results for both M and NM analyses are shown in Figure 5.

GLOBETROTTER was then used to identify and date admixture events. We started with a NM analysis, using ‘prop.ind:1’ and ‘null.ind:1’ options. This allowed us to estimate whether admixture was detected and date the event(s). With ‘prop.ind:0’ and ‘bootstrap.date.ind:1’ options, 100 bootstraps were run estimating the C.I. around the previously calculated date. We used the date estimates ≤1 and ≥400 generations as evidence of admixture, calculating a p-value as in [2]. We thus considered p-values ≥0.01 as support to the null hypothesis of no admixture detected.

Out of 10 targets we tested, 8 gave evidence of admixture and were further tested with GLOBETROTTER, this time using ‘prop.ind:1’ and ‘null.ind:0’ options. In this step we defined the admixture proportions of the sources involved in the admixture event(s) and described them as a mixture of surrogate populations (including the French groups).

For the 8 passing targets we also ran a M analysis, this time masking the French clusters from the group of surrogates. All 8 targets passed the first step (‘null.ind:1’, ‘prop.ind:1). In the following phase (‘null.ind:0’ and ‘prop.ind:1’), one target only was excluded because showing a p-value ≥0.01. In this work, we show the results from the M analysis for 5 out of 7 targets (Supplementary Figure 9), since we only considered those targets whose reconstructed sources were described by at least one surrogate. In one case (target SE), a *multiple-date* was estimated and we changed the result to *one-date-multiway* following the methodology described in ref. [1].

### *References*


1. Busby, G. B. J. *et al.* The Role of Recent Admixture in Forming the Contemporary West Eurasian Genomic Landscape. *Curr. Biol.* **25**, 2518–2526 (2015).
2. Hellenthal, G. *et al.* A genetic atlas of human admixture history. *Science (80-. ).* **343**, 747–751 (2014).
